# Comparative Genomics Reveals LINE-1 Recombination with Diverse RNAs

**DOI:** 10.1101/2025.02.02.635956

**Authors:** Cheuk-Ting Law, Kathleen H. Burns

## Abstract

Long interspersed element-1 (LINE-1, L1) retrotransposons are the most abundant protein-coding transposable elements (TE) in mammalian genomes, and have shaped genome content over 170 million years of evolution. LINE-1 is self-propagating and mobilizes other sequences, including *Alu* elements. Occasionally, LINE-1 forms chimeric insertions with non-coding RNAs and mRNAs. U6 spliceosomal small nuclear RNA/LINE-1 chimeras are best known, though there are no comprehensive catalogs of LINE-1 chimeras. To address this, we developed TiMEstamp, a computational pipeline that leverages multiple sequence alignments (MSA) to estimate the age of LINE-1 insertions and identify candidate chimeric insertions where an adjacent sequence arrives contemporaneously. Candidates were refined by detecting hallmark features of L1 retrotransposition, such as target site duplication (TSD). Applying this pipeline to the human genome, we recovered all known species of LINE-1 chimeras and discovered new chimeric insertions involving small RNAs, *Alu* elements, and mRNA fragments. Some insertions are compatible with known mechanisms, such as RNA ligation. Other structures nominate novel mechanisms, such as trans-splicing. We also see evidence that LINE-1 loci with defunct promoters can acquire regulatory elements from nearby genes to restore retrotransposition activity. These discoveries highlight the recombinatory potential of LINE-1 RNA with implications for genome evolution and TE domestication.

**GRAPHICAL ABSTRACT:** 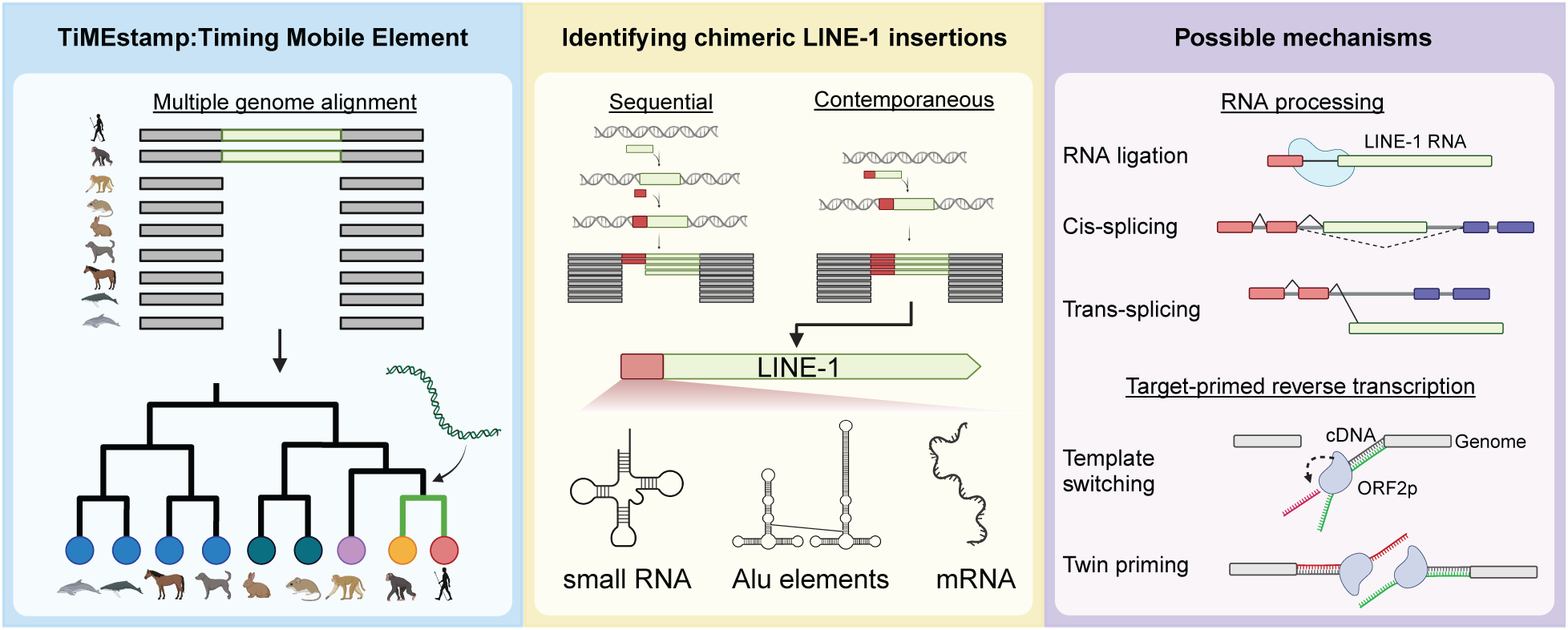

## INTRODUCTION

Mammalian long interspersed element-1 (LINE-1, L1) retrotransposons are non-long terminal repeat (non-LTR) transposable elements (TEs) and represent the most abundant protein-coding, self-propagating retroelements in their hosts (1–4). While LINE-1 protein-coding sequences are relatively conserved, active LINE-1 promoters are variably sized (1–2 kb) and have different sequences across species (1,5). Their persistent activity over more than 170 million years (Myr) of mammalian evolution has profoundly shaped genomes; approximately 30% of the human genome (1 billion base pairs) has been written by the LINE-1 reverse transcriptase (6–8).

Humans inherit a compliment of retrotransposition-competent LINE-1 loci, which are transcribed by the LINE-1 internal CpG rich RNA polymerase II promoter. Its RNA encodes two proteins: open reading frame 1 protein (ORF1p) (9,10), an RNA-binding protein, and ORF2p (11–13), which possesses endonuclease (EN) and reverse transcriptase (RT) activity. During retrotransposition, these activities work in concert to execute target-primed reverse transcription (TPRT), using a genomic DNA nick to prime reverse transcription of the LINE-1 RNA to complementary DNA (cDNA). TPRT intermediates are resolved to yield double stranded LINE-1 genomic insertions (14–18). The process produces several sequence hallmarks that characterize ORF2p mediated insertions, including target-site duplications (TSDs) (19–21), which are short duplicated sequences flanking both ends of the insertion. Another hallmark is the presence of an endonuclease recognition motif (TT/AAAA, slash indicates the cut site) (16,22,23). Lastly, LINE-1 insertions typically have a poly(A) tail at their 3’ end and are variably 5’ truncated (24,25).

Apart from copying its own sequence (in *cis*), LINE-1 ORF2p is responsible for reverse transcribing other sequences into the genome. The most notable examples in humans are non-coding retrotransposons including *Alu* (26) and composite ‘SINE-R, VNTR, and *Alu*’ (SVA) elements (27,28). By hijacking the TPRT mechanism, *Alu* and SVA have generated approximately 1.4 million and 3 thousand copies in the human genome, respectively (6). LINE-1 also drives the insertion of processed (spliced) mRNAs or pseudogenes in *trans* (29,30) and retrotransposes small non-coding RNAs, including transfer RNAs (tRNAs) (31), Y RNAs (32), small nuclear RNAs (snRNAs) (33–35), small nucleolar RNAs (snoRNAs) (36) and ribosomal RNAs (rRNAs) (31) in genomes. Collectively, hundreds of thousands of copies of these non-coding RNA-derived sequences are present in the genome.

LINE-1 is known to produce chimeric insertions with some of these RNA species, fusing short non-LINE-1 sequences with the 3′ end of a LINE-1 in a single insertion event. To date, LINE-1 has been reported to form chimeric insertions with U1-U6 snRNA (33–35) and 5S rRNA (37). Among these, U6/LINE-1 chimeric insertions are most frequent and best characterized; in these chimeras, the U6 segment is consistently full-length, and occurs 5′ of the LINE-1 and in the sense direction relative to the 3′ LINE-1 end (35). The incorporation of the 3′ end of LINE-1 suggests that some mechanisms of *cis* preference likely contribute to insertion (30), and like other TPRT events, these chimeras often occur at the LINE-1 endonuclease motif and are flanked by TSDs. The formation of U6/LINE-1 chimeric insertion may occur through two distinct mechanisms: RNA ligation, generating a chimeric RNA template for reverse transcription (38), or template switching during TPRT, where the ORF2p RT shifts from the its primary LINE-1 RNA template to a nearby RNA molecule, concatenating the cDNA of both (35).

Building a comprehensive census of 3′-LINE-1 chimeras in the genome has been challenging. New chimeras have primarily been identified by comparing locations of a ‘bait sequence’ (a known sequence with a predilection for chimera formation, e.g., U6) with LINE-1 elements to identify adjacent repeats (33–35,37). Candidate chimeric insertions can then be confirmed by annotating TSDs and other sequence signatures of retrotransposition.

Degradation of sequence homologies over time and independently occurring “nested” insertions of more recently active TEs can obscure key sequence signatures and pose challenges for recognizing chimeras. Furthermore, this approach heavily relies on *a priori* knowledge of bait sequences, limiting the discovery of novel of chimeric insertions.

To address these challenges, we developed a computational pipeline - TiMEstamp that applies a comparative genomics approach to estimate the insertion time of LINE-1 and identify chimeric LINE-1 insertions without reliance on bait sequences. First, we identify insertions encompassing LINE-1 and non-LINE-1 sequences which appear contemporaneously in evolutionary history using genome-wide multiple sequence alignment (MSA) of 464 genomes across 438 mammalian species (39). Next, we filter these candidates by searching for sequence hallmarks of LINE-1-mediated insertions. To overcome challenges posed by sequence divergence over time, we derive consensus sequences of insertion alleles using MSA, an approach which greatly improves detection of TSDs.

Using this pipeline with the human genome as a reference, we identified all known species of 5′-L1 chimeras as well as a previously unrecognized set of RNA repeats, *Alu* elements and fragments, and mRNA transcripts that have formed chimeric insertions with LINE-1 over a broad timespan in mammalian evolution. These findings indicate new mechanisms by which LINE-1 forms chimeric insertions, including *trans*-splicing events. Additionally, we see evidence that LINE-1 insertions with defunct promoters can acquire regulatory elements from nearby genes that restore the capacity for retrotransposition. These discoveries underscore the recombinatory potential between cellular RNAs and retroelements with implications for transposon evolution and TE domestication.

## MATERIALS AND METHODS

### Preparing repeat datasets

Repeat annotations for the hg38 genome assembly (version: Repeat Library 20140131) were obtained from www.RepeatMasker.org. Over evolutionary time, LINE-1 sequences can be fragmented by other transposable element insertions (i.e., ‘nesting’). To address this, distinct LINE-1 fragments putatively derived from the same insertion, predicted by RepeatMasker (40), were grouped based on their repeat ID. The orientation of each LINE-1 insertion was determined by the strand of the fragment closest to the 3′ end of the LINE-1 consensus sequence. To predict candidate chimeric LINE-1 insertions, we excluded LINE-1 elements with 5′ inversions by requiring the same orientation across all fragments. The 5′ most end of each LINE-1 insertion was then used for chimeric LINE-1 predictions. In contrast, for evolutionary studies in Figure 2, LINE-1 fragments with the same repeat ID were consolidated into potentially single insertions, regardless of their respective orientations. Additionally, consensus sequences representing each LINE-1 family were retrieved from Khan et al. (2006) (5).

### Preparing multiple sequence alignments

Multiple sequence alignment (MSA) data in multiple alignment format (MAF) format, encompassing 470 mammalian genomes from 444 species, were prepared by Hiller and colleagues as described in Hecker et al (39) and retrieved from the UCSC Genome Browser (https://hgdownload.soe.ucsc.edu/goldenPath/hg38/multiz470way/), together with the phylogeny tree (41). MAF files were split by species, and gaps relative to the human reference genome (hg38) were removed using the Phylogenetic Analysis with Space/Time (PHAST) program commands mafSpeciesSubset -keepFirst and msa_view --gap-strip 1 (42). Since precise alignment within the MSA was not critical for our analysis, we focused on unaligned regions to reduce data size. These unaligned regions, indicated by asterisks (*) or hyphens (-) in the resulting FASTA files, were extracted and annotated in BED file format. Additionally, we excluded non-ape sister clades containing fewer than 10 species, as smaller clades are less effective at reducing noise through averaging, especially when evolutionarily distant from humans. After filtering, 464 genomes, including humans, were retained.

### TiMEstamp: Calling presence/absence of LINE-1 from MSA data and predicting their insertion times

Presence or absence of a LINE-1 insertion across different mammals allowed us to place its arrival time with respect to speciation. Before classifying the presence or absence of LINE-1, in each species, we calculated the aligned portion to annotated LINE-1 in the human genome, regardless all the mismatches and averaged the aligned portion across species within the sister clade. To ensure that absent LINE-1 alignments were restricted to pre-insertion loci and not caused by a larger genomic deletion or absence encompassing the LINE-1, regions with upstream gaps exceeding 1000 bp were classified as ‘missing’ the insertion site. All species in a non-ape sister clade were marked as missing an insertion site if the site was missing from 90% of clade members. For evolutionary analyses in figure 2, we labeled an insertion as “present” in a sister clade if the average aligned portion exceeds 50% of annotated LINE-1. For chimeric LINE-1 analysis, we used a threshold of 65% alignment (two-thirds) to classify a segment as “present”, while no or shorter homologous intervals were classified as “absent”. Using the presence or absence of the LINE-1 sequence in sister clades, the earliest observed presence was used to infer the timing of its initial introduction.

### Defining insertions 5′ of LINE-1 and predicting their insertion times

Sequence insertions 5′ of LINE-1 but lacking homology to LINE-1 were found by identifying ‘gaps’ (relative to the human genome) between LINE-1 insertions annotated in the human genome and upstream sequences present in genomes lacking the LINE-1 insertion. Gaps shorter than 1000 bp and longer than 25bp were included and were aggregated together if they were within 20 bp of each other. These gap clusters had to be present in at least 15 species. For each gap cluster, the gap supported by the most species (with a minimum of 5 species) was selected as the representative gap. Among the representative gaps, the shortest one was designated as the representative upstream region of LINE-1. The timing of the sequence’s introduction was inferred using the same method applied for LINE-1, but with a stricter length of homology threshold of 75%. Regions with an average alignment below 75% were classified as absent.

### Predicting chimeric LINE-1 insertions

To identify potential chimeric LINE-1 insertions, the predicted arrival time of the LINE-1 element and its 5′ adjacent insertion were compared. Where the two appear contemporaneous (i.e., are distributed identically within clades and therefore inferred to arrive within the same time interval), the insertions were classified as a candidate chimeric LINE-1 insertion. These were filtered for alignment noise by excluding instances where more than 10% of genomes in the non-ape sister clades without the LINE-1 element, exhibited at least 25% alignment for LINE-1 sequence. MSAs for all presented data were manually inspected, using either the 470-way alignments generated by MULTIZ from Hiller’s lab or the 447-way mammalian MSA produced by Cactus from UCSC Genome Institute and Farh’s lab or both (39,41,43).

### Annotation of the unannotated sequence of the chimeric LINE-1 insertions

To discern the origins of sequences 5′ adjacent to chimeric LINE-1, these upstream sequences subjected to BLAST (44) searches against RNAcentral (45), Gencode (version 46) (46), and Rfam databases (47). The best alignment was determined by considering the aligned length, coverage of the query sequence, and pairwise alignment identity. Alignments exceeding 75 bp and covering >75% of the query sequence were prioritized as representative alignments. In cases where multiple alignments or none met these criteria, the alignment with the highest query coverage was selected. If multiple alignments exhibited identical coverage, the one with the highest pairwise alignment identity was chosen. When several alignments fulfilled all three conditions, the top listed alignment was selected as the final representative. For chimeric insertions fusing protein-coding mRNAs or long non-coding RNAs (lncRNAs) with LINE-1, annotations were also manually verified.

### Building a census of known chimeric LINE-1 insertions

To compile a census of chimeric LINE-1 insertions with known structures (e.g., U1-6 small nuclear RNA fused to 5′ LINE-1 and 5S ribosomal RNA fused to 5′ LINE-1), we identified LINE-1 where these short RNA species were annotated by RepeatMasker within 50 bp upstream of the LINE-1 element. The identified loci were manually inspected to evaluate the quality and clarity of their MSA profiles to confirm their contemporaneous insertion, ensuring the accuracy and reliability of the curated dataset.

### Determining target site duplications (TSDs)

Flanking TSDs offer molecular evidence that compound or complex insertions occurred together in a single target primed reverse transcription (TPRT) event. To delineate TSDs at insertion junctions while permitting for some single nucleotide substitutions since the insertion, sequences flanking insertions were extracted and converted into IUPAC ambiguity codes. For TSD prediction flanking LINE-1 elements, sequences extracted by bedtools (48) encompassed 75 bp upstream and 25 bp downstream of the 5′ junction, and 25 bp upstream and 75 bp downstream of the 3′ junction. The IUPAC-converted sequences were aligned using a Smith– Waterman local alignment algorithm (49) to detect regions of similarity at the junctions. A modified scoring matrix, adapted from the DNAfull scoring matrix (50), was employed. In this scoring matrix, we reduced the match score was from 5 to 3, and increased the mismatch penalty from −4 to −10, effectively suppressing the extension of alignments beyond mismatched regions. Additionally, the gap opening penalty was set to −50 to minimize the occurrence of gaps in the alignments. For LINE-1, predicted TSDs were considered high-quality if their lengths ranged from 10 to 30 bp and the end of the TSD at the 5′ junction was located within 3 bp upstream or downstream of the junction of the LINE-1 element. For chimeric LINE-1 insertions, the TSD length requirement was adjusted to 8 to 20 bp, and the end of the TSD at the 5′ junction was allowed to occur within a broader range, between 40 bp upstream and 25 bp downstream of the predicted 5′ junction. In cases where the 3′ end of the LINE-1 was poorly annotated due to polyA homopolymers in the preinsertion sequence, TSDs were manually curated.

### ORF2p endonuclease (EN) motif analysis

For the ORF2p EN motif analysis of the L1MB4 family, we extracted sequences spanning the 5′ junction of the TSD at the 5′ end of LINE-1, including 4 bp upstream and downstream of the junction. These sequences were then analyzed for motif enrichment using the MEME Suite (51). For chimeric insertions, the two base pairs immediately upstream of the TSD at the 5′ end of LINE-1 were identified as potential ORF2p EN motifs.

### Transposable Element Age Analysis

The age of transposable elements (TEs) was calculated from the percentage divergence values provided by RepeatMasker. These values represent the percent substitution of each TE from its consensus sequence. Using the Jukes-Cantor model (52), the divergence was corrected for multiple substitutions, and the age was estimated in millions of years (Myr) based on a substitution rate of 0.17% per million years (5).

### Transcriptomic data analysis

Oxford Nanopore Technology (ONT) long-read sequencing data were downloaded from the NIH Sequencing Read Archive (SRA) (Accession number: PRJEB81685) (53). Sequencing reads were aligned to the *RAP1GDS1* segment of the *RAP1GDS1*/LINE-1 chimeric insertion using minimap2 (54). Extracted sequences were then aligned to the L1PA2 sequence to identify chimeric transcripts. The expression of *RAP1GDS1* in primates was retrieved from a precomputed gene expression matrix in the Gene Expression Omnibus (GEO) (Accession number: GSE127898) (55).

### RNA secondary structure prediction

5S rRNA, tRNA and 28S rRNA (LSU-rRNA) Sequences were extracted from Dfam (56) and analyzed using RNA Secondary Structure Determination Tool (R2DT) (57), a tool integrated with RNAcentral, to predict and visualize RNA secondary structures.

## RESULS

### Inferring the Occurrence of a Retrotransposon Insertion Using Multiple Sequence Alignment (MSA)

Retrotransposon insertions are mutagenic events which are homoplasy-free and which have unambiguous directionality. In other words, insertion of a specific retroelement sequence at a specific site in the genome with associated changes to the target site collectively represent unique features of a single mutagenic event. Seeing the same features in a different genome essentially always reflects identity by descent and never two independently occurring insertions. Further, the ancestral or antedating allele is always the ‘preinsertion’ allele; the derived allele is always the insertion allele. Thus, comparative genomics using multiple sequence alignment (MSA) across species is a powerful tool to delineate the structure of retrotransposon insertions and to infer the timing of the insertion with respect to speciation.

We focused on studying the retrotransposon composition of the human genome, with a particular emphasis on LINE-1 elements, which represent one of the most abundant transposable element families, with over 500,000 copies identified (6). LINE-1 elements can be divided into three major types based on evolutionary age: the oldest, L1M (including L1MA-E families, which are mammalian-specific); L1P (including L1PA and L1PB, which are primate-specific); and the youngest, L1HS (also referred to as L1PA1, which is human-specific). Based on sequence homology studies, the L1M families are estimated to have been active approximately 60–150 million years ago (Mya), the L1P families between 7–80 Mya, and L1HS around 3 Mya (5,58,59). This prolonged activity has resulted in the significant accumulation of LINE-1 elements in the human genome.

To infer the insertion timing of mobile elements with respect to speciation, we developed a computational pipeline – TiMEstamp to analyze a publicly available MSA of 464 genomes from 438 species, computed by Hiller’s group and accessible via the UCSC Genome Browser. The MSA was generated by pairwise aligning sequences to the human genome using the LASTZ aligner (60), followed by constructing the MSA with MULTIZ (61), as detailed in a previous study (39). We transformed the resulting alignment into a binary format, where regions were classified as either aligned (indicating the presence of the target sequence) or gapped (indicating its absence), without requiring nucleotide-level agreement. An alignment score of 1 indicated the sequence was fully present, while a score of 0 indicated its complete absence. Scores between 0 and 1 may be attributable to deterioration of sequence homology over time or artifacts of alignment (Supplementary Figure 1). To minimize the noise from a particular genome, we averaged alignment scores at the sister clade level, where each clade shares a single common ancestor diverged from the human lineage at the same branchpoint (Figure 1). Using these scores, we next classified insertions as present or absent within each clade. We concluded by placing the time of arrival of each insertion at the common ancestor of a monophyletic clade where the insertion is present in all members of that clade and absent in other taxa.

**Figure 1.**
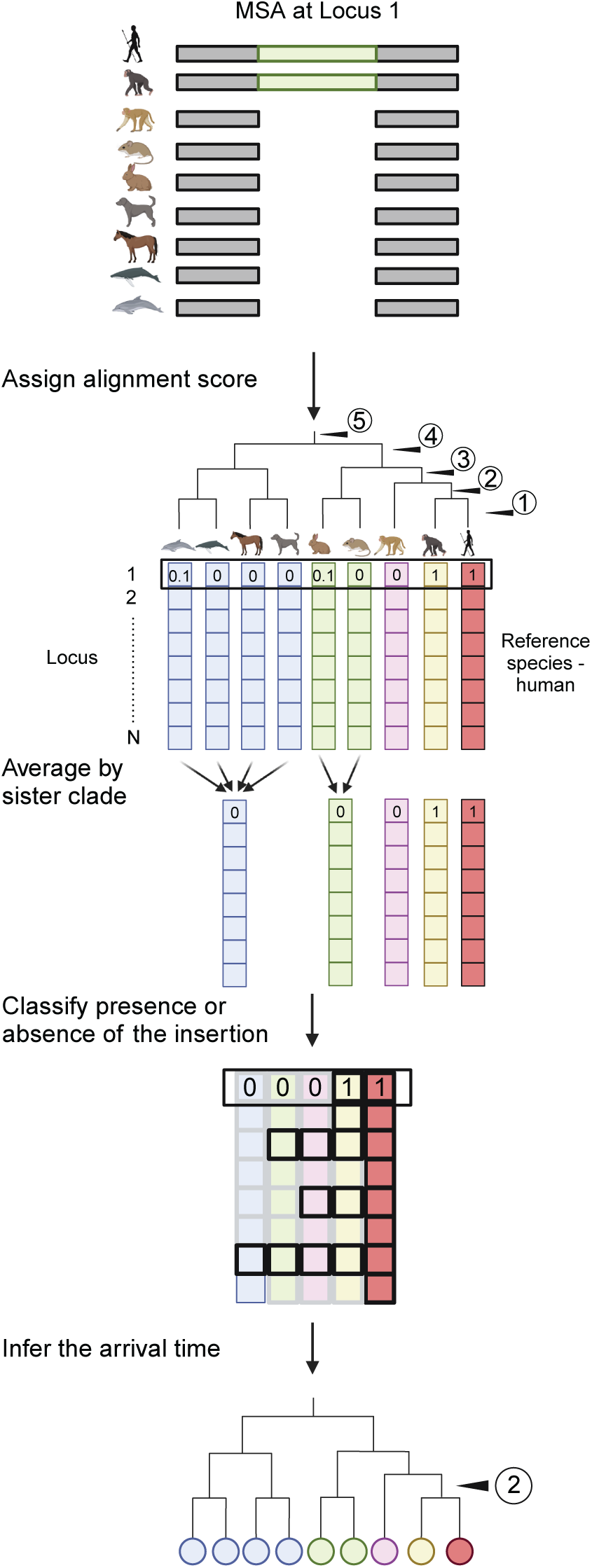
TiMEstamp: A computational pipeline for inferring the evolutionary timing of sequence insertions. Multiple sequence alignment (MSA) data are used relative to the human genome to define ancestral (pre-insertion) vs. insertion alleles. In each species, alignment to an insertion found in the human genome is scored 1 (complete alignment), signifying shared presence of the sequence; 0 (no alignment) representing a gap or absence of the sequence; or an intermediate value. Scores between 0 and 1 may be attributable to deterioration of sequence homology over time or artifacts of alignment. To reduce noise, alignment scores are averaged at the sister-clade level stepping away from the human reference. An insertion is inferred to have originated in a common ancestor of a monophyletic clade when it is present in all members of that clade/taxon and absent in other taxa/taxon.

**Figure 2.**
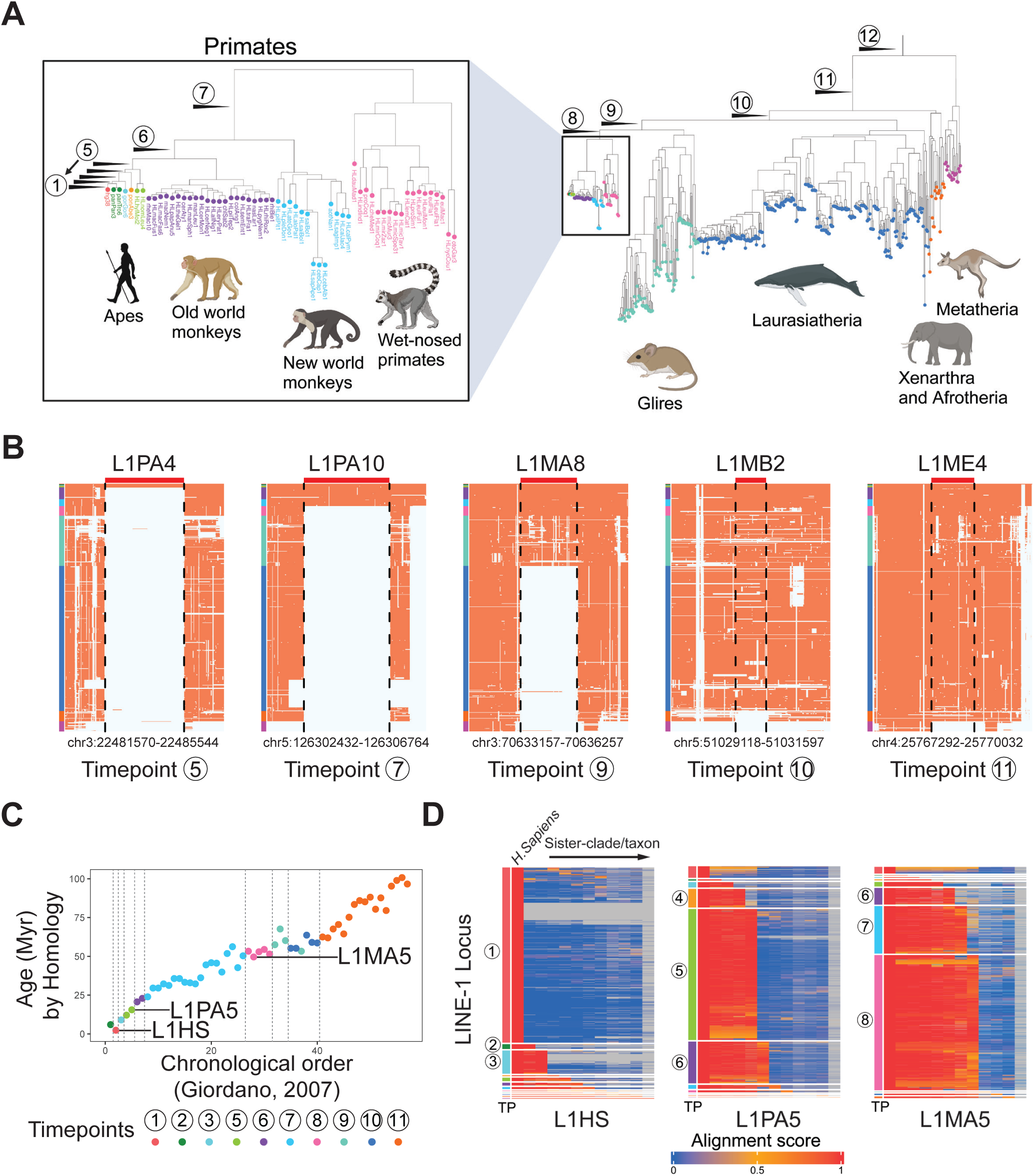
Inferring LINE-1 insertion times in mammalian evolution. (A) The phylogenetic tree of 464 genomes across 438 species organizes into 11 sister clades/taxon and *Homo sapiens* (highlighted in different colors) and 12 ancestral time points. Species having more than one genome in the dataset are mostly from the Laurasiatheria clade. (B) MSA profiles of 5 LINE-1 loci. Chromosomal locations are from the human genome and alignments of syntenic intervals in other species are arranged vertically. The color labels on the far-left side of each profile corresponds to the sister clade from panel A; red in the MSA represents alignment, while grey indicates gaps or no alignment. LINE-1 subfamilies (L1PA4, L1PA10, L1MA8, L1MB2, and L1ME4) were annotated by RepeatMasker; inferred time points of each insertion are noted below the MSA profile. The genomic coordinates underneath each heatmap refer to genomic location of the corresponding LINE-1 locus. (C) Inferred insertion timepoints (dot colors), the chronological order of LINE-1 subfamilies calculated from defragmentation (X-axis) and ages, million year (Myr) calculated using divergence from the subfamily consensus sequence (Y-axis) are highly correlated. No LINE-1 subfamily was classified as inserting at Timepoint 4. L1PA4 activity spans both Timepoints 4 and 5; its classification was attributed to Timepoint 5 due to a slightly higher number of loci assigned to that timepoint. (D) Alignment scores against the human genome for three LINE-1 subfamilies (L1HS, L1PA5, and L1MA5). Each row represents a distinct locus in the subfamily; each column corresponds to the sister clade/taxon-level averaged alignment scores(far left column: *Homo Sapiens*, far right column: Metatheria). The inferred insertion timepoint (TP) for each locus is indicated on the left of the graph. Most L1HS inserted at timepoint 1; 244 loci (18.1%) appear to have earlier insertion timepoints. Most L1PA5 inserted at timepoints 5 and 6, while L1MA5 primarily integrated at timepoints 7 and 8. This visualization highlights the successive activity intervals of these LINE-1 families.

### Inferring the arrival of LINE-1 families in the human lineage

Phylogenetic analyses of the 464 species shows that these group as 11 sister clades (or taxons) with respect to the human reference, defining 12 distinct timepoints when we can infer LINE-1 elements inserted into the ancestral human genome (Figure 2A). We can appreciate, for example, that the primate specific L1PA4 subfamily (Figure 2B) appeared subsequent to the divergence of apes and monkeys (corresponding to timepoint 5 in Figure 2A), while L1PA10 (Figure 2B) was introduced earlier, after the split between New World and Old World monkeys (at timepoint 7, Figure 2A). Mammalian L1M subfamilies antedate these. L1MA8 was present in the genomes of a common ancestor shared by primates and Glires (e.g., rodents and rabbits). In contrast, older LINE-1 elements, such as L1MB2, were shared in all species evaluated except Xenarthra and Afrotheria (e.g., moles and elephant) and Metatheria (e.g., kangaroo) clades, or in the case of L1ME4, absent only in Metatheria (Figure 2B).

To evaluate the accuracy of our approach, we tested these inferences against RepeatMasker annotations (40). To establish that standard, Smit et al. used 3′ sequence homology to infer the history of LINE-1 elements and annotate LINE-1 families (59). Using our approach, which incorporates MSA data, we confirmed that L1P families are strictly primate-specific. Consistent with the original finding and other studies (5,58), the youngest L1M families, L1MA3 through L1MA1, are also primate-specific, whereas L1MA6 through L1MA4 were active during the divergence of primates from Gilires and continued to be active following the emergence of primates (Supplementary table 1).

Furthermore, as expected, this ordering closely aligns with LINE-1 ages calculated by homology to subfamily consensus sequences and orders reconstructed by defragmentation analyses (58) (Figure 2C). As examples, we illustrate sister clade-based alignment scores for all loci of three representative LINE-1 subfamilies: the *Homo sapiens* associated L1HS, the primate LINE-1 subfamily L1PA5, and the mammalian LINE-1 subfamily L1MA5 (Figure 2D). In each map, rows represent individual loci within the subfamily, while columns display the sister clade-level averaged alignment scores. The inferred insertion timepoint for each locus is shown on the left side of the graph. As expected, most annotated L1HS loci are human-specific; however, 176 loci (13.1% of total loci annotated as L1HS in RepatMasker) are shared with chimpanzee (62) and gorilla (63) and thus classified as having earlier insertion times. This finding suggests that these loci would be better labeled as primate LINE-1. Notably, approximately 90% of these loci lack the 3′ UTR sequence and the 3′ 700bp of ORF2, key features used for LINE-1 family classification (59), and so these are challenging loci for RepeatMasker to accurately assign. For L1PA5, most loci were introduced at timepoint 5, corresponding to the divergence between apes and Old World monkeys. L1MA5 displayed a broader activity interval, spanning timepoints 7 and 8, dating back to after the split between primates and Glires. Collectively, these findings confirm the utility of MSA to recapitulate the known succession of LINE-1 activity over evolutionary time.

### Distinguishing Sequential and Chimeric LINE-1 Insertions Using MSA

MSA provides insight not only into when a DNA segment was introduced into the genome but also into whether two adjacent DNA segments were inserted sequentially or contemporaneously. Sequential insertions are inferred when one DNA segment is found in older evolutionary groups, while the neighboring sequence appears only in more recently evolved groups. This pattern indicates that the genomic integrations occurred at different times, one following the other. On the other hand, contemporaneous insertions are revealed when both DNA segments are consistently present together across all evolutionary groups in the MSA. This pattern indicates that the segments were inserted around the same time, potentially integrating in a single event (Figure 3A).

**Figure 3.**
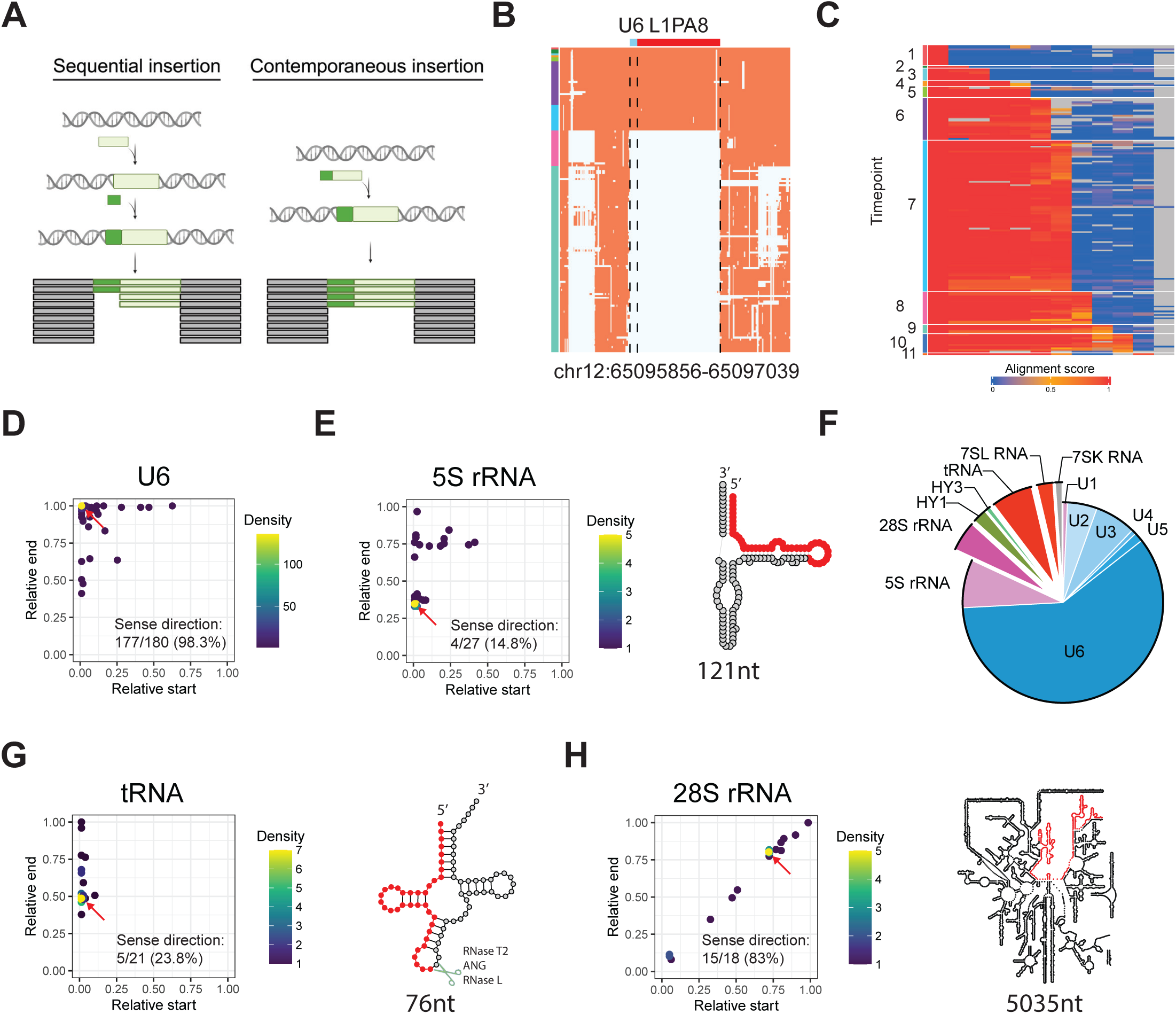
Characterization RNA repeat/LINE-1 chimeras. (A) MSA data reveal the arrival times of two adjacent DNA segments, enabling a distinction between sequential and contemporaneous insertions. (B) The MSA profile of a chimeric U6/LINE-1 insertion illustrates how the L1PA8 and U6 were inserted into the genome contemporaneously. (C) The heatmap shows the human and sister-clade/taxon level alignment score (columns) for all 180 identified chimeric U6/LINE-1 loci (rows). This pattern suggests production of chimeric U6/LINE-1 insertions has spanned a long timeframe of LINE-1 activity. (D) The scatter plot shows the relative start (X axis) and end (Y axis) of U6 segments, normalized by the reference U6 length. U6 segments in the majority of chimeric U6/LINE-1 insertions are full-length and in the sense orientation relative to LINE-1. The red arrow points to the dot that has the highest density of data points. (E) The 5S segment in chimeric 5S/LINE-1 insertions is often truncated and oriented in the antisense direction relative to LINE-1. On the right side, the predicted RNA secondary structure of 5S rRNA is shown, with the region frequently incorporated into chimeras highlighted in red (1-45nt). (F) The pie chart shows the relative abundance of every RNA repeat identified in chimeric insertions with LINE-1. Newly identified chimeras, highlighted in the enlarged section, include 28S rRNA, HY1, HY3, tRNA, 7SK RNA and 7SL RNA. Others were previously known RNA repeats associated with chimeric LINE-1 insertions. RNA groups are color-coded by their RepeatMasker annotation: blue, small nuclear RNA (snRNA); grey, signal recognition particle RNA (srpRNA); purple, ribosomal RNA (rRNA); green, small cytoplasmic RNAs (scRNA); and red, transfer RNA (tRNA). (G) The scatter plot shows the relative start and end sites of tRNA segments in tRNA/LINE-1 chimeras, normalized by the reference tRNA length. Most tRNAs in tRNA/LINE-1 chimeras are half-length and in an antisense orientation relative to the LINE-1. On the right side, the RNA secondary structure of tRNA is shown, with the region frequently incorporated into chimeras highlighted in red (1-37nt). tRNA can be cleaved by ribonucleases, such as RNase T2, angiogenin (ANG), and RNase L, to generate tRNA halves. (H) The scatter plot shows the relative start and end sites of 28S rRNA segments in 28S rRNA/LINE-1 chimeric insertions, normalized by the full-length 28S rRNA. Most 28S rRNA segments are in sense orientation relative to LINE-1. On the right side, the RNA secondary structure of 28S rRNA is shown, with the region frequently incorporated into chimeras highlighted in red (3636-4115nt).

This approach allows us to identify potential chimeric LINE-1 insertions. Chimeric LINE-1 insertions occur when a LINE-1 retrotransposon integrates along with additional, unrelated sequences at its 5′ end in a single mutagenic event. U1, U2, U4, U5, U6 snRNA, U3 snoRNA and 5S rRNAs are all known to form chimeric insertions with LINE-1 (34,35,37,64). To begin, we compiled a catalogue of chimeric insertions of these types by manually inspecting MSA profiles of LINE-1 elements that have these RNA sequences nearby. All together, we identified 253 candidate chimeric insertions (Supplementary Table 2), the most comprehensive catalog to date. An example MSA of a U6/LINE-1 chimeric insertion is shown in Figure 3B. Among these chimeras, 70.9% involved recombination between LINE-1 and U6 spliceosomal snRNA, 10.5% involved 5S rRNA, and 8.5% involved U3 snoRNA. Their relative abundance is consistent with prior studies (34,35,37,38,64).

Insertions of U6/LINE-1 chimeras have occurred recurrently in human evolution; each appears to be the product of a unique recombination event joining a U6 and the 3′ end of a LINE-1, and these do not continue to self-propagate after insertion. The majority have accumulated between timepoint 8, following the divergence of primates and Glires (approximately 90 million years ago), and timepoint 6, after the separation of monkeys and apes (around 30 million years ago) (Figures 2A, 3C). In these chimeras, the U6 portion is often full-length (106 nucleotides), while the LINE-1 portion is frequently 5′ truncated (33,35,37).

Interestingly, full-length U6 sequences are more abundant in U6/LINE-1 chimeric insertions compared to isolated U6 in the genome (Supplementary Figure 2A). Meanwhile, the LINE-1 5′ truncation in these chimeras is more pronounced than considering the length distribution of all genomic LINE-1 (Supplementary Figure 2B). Consistent with prior studies, almost all chimeric U6 (98.3%) are oriented in the same direction as their associated LINE-1 (sense orientation) (Figure 3D). These recurrent patterns indicate a common mechanism underlies these chimeric insertions, potentially an RNA ligation creating a contiguous U6-3′ LINE-1 template before TPRT (38) or a LINE-1 RT template switch concatenating these sequences. Similarly, U3 sequences in U3/LINE-1 chimeric insertions exhibit a strong bias toward the sense orientation (95.5%). However, they are predominantly found in a truncated form (Supplementary Figure 2C), contrasting with the bimodal length distribution of other genomic U3 sequences (Supplementary Figure 2D). This suggests that U3 fragments are more likely than full-length U3 snoRNAs to recombine with LINE-1 or that U3 snoRNA sequence may be lost during the recombination and/or integration to form the chimeric insertion.

5S rRNA segments in 5S/LINE-1 chimeric insertions are frequently oriented in antisense with respect to their associated LINE-1 and are truncated. Typically, the start of the 5S rRNA is at the junction with the LINE-1 and extends until its truncation at position of 46 and 90 (Figure 3E). These two sites correspond to the Loop C and D of the 5S rRNA. The frequency of truncation at position 46 is more frequent that the total 5S rRNA copies in the genome (Supplementary Figure 2E). Of 8 randomly selected 5S/LINE-1 chimeric loci, 6 show 7-15bp TSDs flanking the insertion and 4 occur at the LINE-1 EN motif (Supplementary Table 2). All U5 sequences observed in U5/LINE-1 chimeric insertions were truncated at their 3′ ends, and 80% found in the antisense orientation relative to LINE-1.

These observed characteristics of U3, U5, and U6 chimeric LINE-1 insertions are consistent with previous reports. Given our larger catalog of events, however, we are better positioned to see recurrent pattens for less common LINE-1 chimeras (e.g., length distributions, RNA repeat breakpoints, and orientation tendencies), and to develop and test hypotheses for how these insertions arise.

### Novel RNA species that form chimeric LINE-1 insertions

To discover new types of chimeric LINE-1 insertions, we developed a computational module in TiMEstamp to systematically identify sequences immediately 5′ (upstream) of LINE-1 fragments inserted contemporaneously with the downstream LINE-1 by MSA. The pipeline delineates these upstream sequences as a gap not aligning to the pre-insertion allele. These sequences are extracted and then compared to existing annotations in the RepeatMasker database (40), transcript annotation of Gencode annotation (46), long non-coding RNA (lncRNA) database, RNAcentral (45) and human genomic sequence (hg38) (6).

Using this pipeline, we successfully identified 76% of the previously manually curated chimeric LINE-1 insertions. Unrecovered loci were predominantly filtered out owing to alignment artifacts. Others were missed because of regional deletions in a sister clade that confounded our inference of insertion timing (Supplementary Figure 3). Beyond the species of LINE-1 chimeras already described, we uncovered five additional types of RNA that contribute to chimeric LINE-1 insertions: tRNA, 28S rRNA (or LSU-rRNA), 7SL RNA, Y RNA (HY1 and HY4), and 7SK RNA (Figure 3F). These newly identified chimeric insertions are significantly less abundant than the previously known examples, with occurrences ranging from a few instances to a maximum of 21 loci.

Among these new types of insertions, chimeric LINE-1 involving tRNA were the most common, with 21 loci detected (Supplementary Table 3). We found 9-19bp TSDs in all 8 randomly selected cases and an EN motif at the insertion site for 6 of them, providing strong molecular evidence that these were inserted together in a single retrotransposition event. Interestingly, the tRNA sequences in these chimeric insertions are typically truncated, comprising approximately half the total length of a full tRNA molecule (Figure 3G). Genomic tRNAs are more commonly intact, 53.4% full-length as annotated in RepeatMasker (Supplementary Figure 2F). Recent studies have shown that tRNAs are frequently cleaved into smaller fragments, known as tRNA halves (30-40nt), by specific RNases, such as RNase T2 (65,66), Angiogenin (ANG) (67), and RNase L (68) (Figure 3G). Breakpoints of the tRNAs in chimeric LINE-1 insertions closely resemble these cleavage sites. This suggests that the tRNA portions incorporated into chimeric LINE-1 insertions may originate from these tRNA halves.

Like 5S rRNA in chimeric LINE-1 insertions, the tRNA sequences are predominantly oriented in the antisense direction relative to the LINE-1 counterpart. This suggests that these chimeras may result from twin-priming, where two reverse transcriptions occur: one at the primary site of 3′ LINE-1 integration and another initiated from the opposite end of a TPRT-induced double-stranded break (69–72).

We discovered a second commonly occurring group of chimeric insertions, 28S rRNA/LINE-1 sequences (n = 18). 28S is a ribosomal RNA, with a full length of 5,035 nucleotides. Among all RNA species associated with chimeric LINE-1 insertions, 28S rRNA segments are the largest, with a median size of 269 nucleotides and a range spanning from 50 to 482 nucleotides. These fragments of 28S rRNA often start around position 3636 in the consensus 28S rRNA sequence (Figure 3H), a feature less commonly seen in other (non-LINE-1) 28S rRNA loci (Supplementary Figure 2G), although segments all along the length of 28S were seen incorporated in 28S/LINE-1 chimeras. Most (83%) of the 28S segments are oriented in the same direction as their associated LINE-1 elements, suggesting that mechanisms such as RNA ligation or template switching during TPRT may underlie their formation.

In summary, our approach identified numerous new RNA repeats that form chimeric insertions with LINE-1. Many of these have recurrent sequence features predicting molecular mechanisms of recombination with LINE-1.

### Using consensus sequences from MSA to detect TSD

To verify each candidate chimeric LINE-1 insertion, we looked for sequence evidence of a single retrotransposition event. Specifically, presence of 10–30 bp TSDs flanking both components of the chimeric insertion supports a model where the intervening sequences result from a single LINE-1 ORF2p-mediated insertion (70,73,74). We tested our ability to identify TSDs by searching for the longest shared sequence at the edges of LINE-1 insertions in the hg38 reference genome. For L1HS, the youngest and most active LINE-1 subfamily in the human genome, we predict TSDs directly from the hg38 genome. Consistent with prior work (74), predicted TSD lengths exhibited a bimodal distribution: one at lengths shorter than 10 bp and another at the range of 10–30 bp, the typical size of TSDs (Supplementary Figure 4A). We speculated that the shorter TSDs likely reflected ambiguous boundaries of LINE-1 sequences or other artifacts. To further evaluate, we analyzed the distance between the predicted TSD (10-30bp) and the 5′ end of the LINE-1, which corresponds to the expected TSD edge. Most TSDs of L1HS are located correctly at the 5′ end (Supplementary Figure 4A). In contrast, for the older LINE-1 subfamily L1MB, fewer TSDs fell within the 10–30 bp range or had TSDs perfectly juxtaposed to the 5′ end of the LINE-1 (Supplementary Figure 4B). We postulated that these discrepancies likely reflected problems with TSD identification attributable to nucleotide substitutions accumulated over time.

To improve TSD identification for relatively older LINE-1 insertions, we leveraged MSA data. We extracted LINE-1 flanking sequences from the hg38 human reference genome and the corresponding sequences in all other species containing the LINE-1 insertions. Ambiguities in the cross-species alignment were resolved using IUPAC codes, enabling both flexibility and precision in sequence alignment (Supplementary Figure 4C). TSDs were identified based on the consensus insertion allele. To test this approach, we analyzed TSDs associated with older LINE-1 families, such as L1MB, TSD predictions based on MSA significantly outperformed those derived solely from hg38. This improvement was evident in the increased number of TSDs with length of 10-30bp and aligning precisely at the 5′ junction of L1MB elements (Supplementary Figure 4D), underscoring the utility of MSA in enhancing the accuracy of TSD predictions for ancient LINE-1 insertions.

Using this method, we benchmarked our computational pipeline across different LINE-1 subfamilies, ranging from the youngest to the oldest, by evaluating the percentage change of TSDs, the proportion with lengths of 10-30bp and the fractions abutting the 5′ junction of LINE-1 elements (+/- 3bp). We see that the MSA-based approach provides incremental improvements in TSD accuracy as the age of the LINE-1 subfamilies increase. The improvement was particularly pronounced for older LINE-1 elements, where TSD predictions using MSA were significantly more reliable than those based solely on the hg38 genome. However, the improvement diminished for very old LINE-1 families, likely due to the reduced quality of MSAs for these ancient elements (Supplementary Figure 4E). Building on this, corrected TSD annotations also improved identification of ORF2p EN motifs associated with insertions (Supplementary Figure 4F). Overall, we conclude that the MSA-based computational pipeline offers a robust and scalable approach for identifying high-quality TSDs, refining EN motif recognition, and providing molecular evidence for ancient LINE-1 insertion events.

### *Alu* forms chimeric insertions with LINE-1

Beyond short RNA species, we identified that *Alu* retrotransposons recurrently form chimeric insertions with LINE-1. To evaluate, we analyzed several candidate chimeric insertions for hallmarks of LINE-1-mediated retrotransposition, namely, TSDs, poly(A)tails, and the ORF2p EN motif (Figure 4A). In total, we identified 113 *Alu*/LINE-1 chimeric insertions (Supplementary table 3).

**Figure 4.**
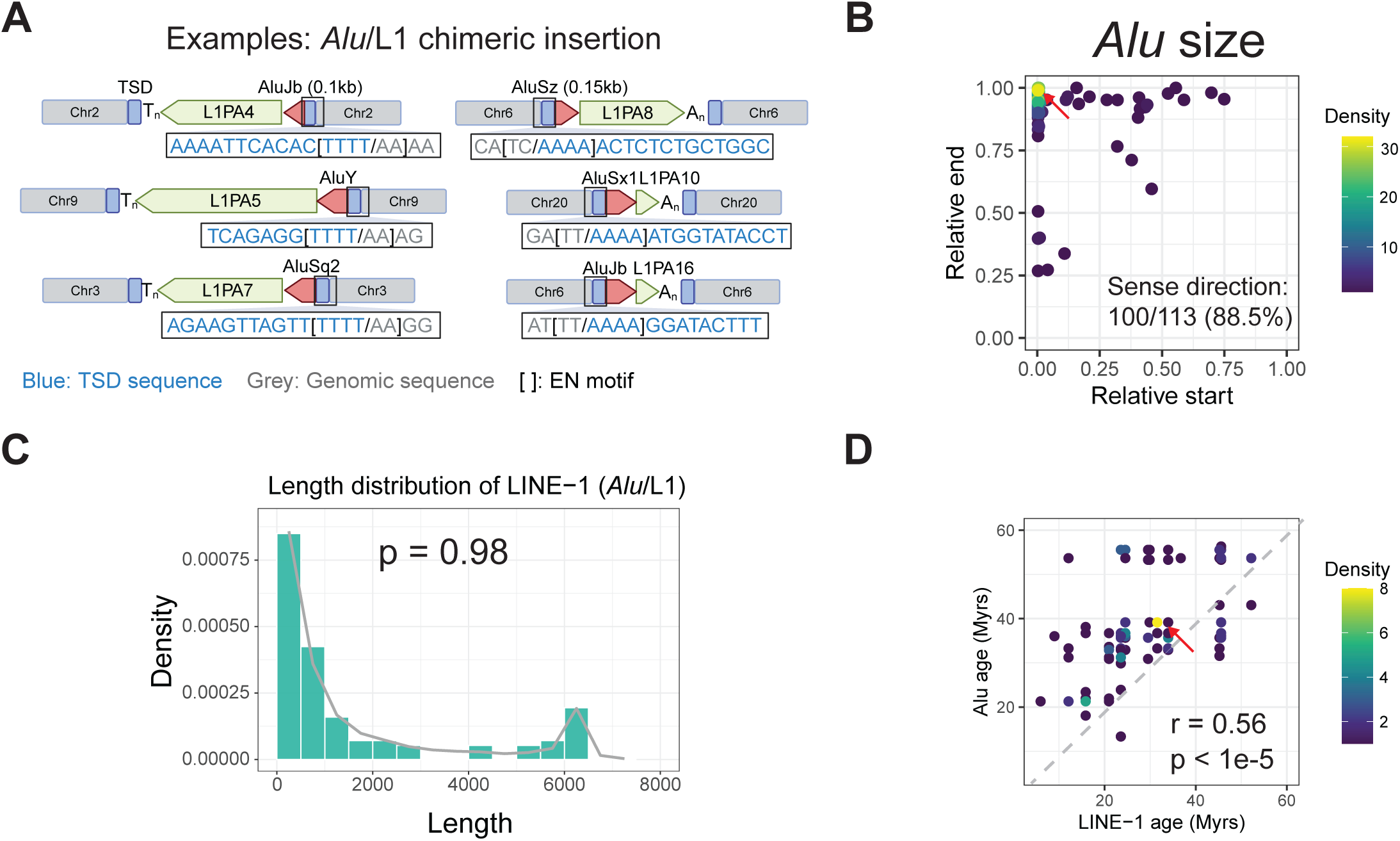
Chimeric *Alu*/LINE-1 Insertions. (A) Examples of *Alu*/LINE-1 chimeric insertions, verified by the molecular signatures of ORF2p mediated retrotransposition, including 11-17 bp of target site duplication (TSD, highlighted in blue), an ORF2p endonuclease (EN) signature motif (shown in brackets: TT/AAAA for sense orientation or AA/TTTT for antisense orientation), and a polyA tail (sense) or polyT tail (antisense). (B) Of the 113 identified *Alu*/LINE-1 chimeric insertions, 88.5% are in the sense orientation relative to LINE-1, with most being full-length insertions. (C) The length distribution of LINE-1 segments in *Alu*/LINE-1 chimeric insertions (green bars) mirrors L1PA length distributions in the genome (grey line). The differences were statistically analyzed using the Wilcoxon’s Rank Sum test. (D) For *Alu*/LINE-1 chimeras, the estimated ages of each LINE-1 component (X-axis) correlates with its paired *Alu*, suggesting chimeras between concurrently active elements. Spearman’s rank correlation coefficient is denoted by r.

To further characterize these chimeric insertions, we examined the length of the *Alu* sequences and their relative orientation with respect to the LINE-1. Most *Alu* elements were full-length (measuring approximately 300 bp) and oriented in the sense direction relative to LINE-1 (88.5%, Figure 4B). Interestingly, the length distribution of the LINE-1 segments in *Alu*/LINE-1 chimeric insertions closely mirrored the overall length distribution of LINE-1 elements in the genome (Figure 4C). We next tested whether component *Alu* and LINE-1 elements in these chimeric insertions were concurrently active and therefore now comparably aged in the genome as assessed by divergence from a consensus sequence. We found a positive correlation between *Alu* and LINE-1 ages for pairs of elements occurring together in a chimeric insertion. We see some tendency for younger LINE-1 elements to form chimeric insertions with older *Alu* families; in contrast, very few instances were observed where a predicted older LINE-1 formed a chimeric insertion with a younger *Alu* element. This asymmetry is consistent with the protein-coding LINE-1 retrotransposon being the primary driver of *Alu*/LINE-1 chimeric insertions (Figure 4D).

Together, these findings indicate that LINE-1-mediated retrotransposition of *Alu* does not always break *cis* preference (30,75), but can arise through the formation of chimeric insertions.

### 5′ Transductions Events forming LINE-1 chimeras with mRNAs and lncRNAs

Using our MSA pipeline, we recognized a second class of 5′ transduction events forming LINE-1 chimeras with unique (non-repetitive) RNA sequences. In total, we identified 17 LINE-1 insertions in the hg38 reference assembly containing RNA sequences fused to the 5′ end of LINE-1, including protein-coding gene messenger RNAs (mRNAs) and lncRNAs (Supplementary table 4). As an example, we highlight a chimeric insertion involving a full-length L1PA5 element and 153bp of upstream sequence (Figure 5A-C). This insertion exhibits all the molecular signatures of LINE-1-mediated TPRT, including a 15 bp TSD flanking both the 3′ end of the LINE-1 and the 5′ transduced sequence, an ORF2p EN motif sequence, and a poly(A)tail at the 3′ end of the LINE-1 segment (Figure 5B). BLAST analysis revealed that the 5′ non-repetitive sequence maps to the first two exons of an isoform transcript of the mitogen-activated protein kinase kinase kinase 13 (*MAP3K13*) gene in the sense orientation (Figure 5C). Further inspection of the genome did not reveal any full-length L1PA5 element in the introns of *MAP3K13*. While such a LINE-1 variant may have existed in an ancestral population, the absence of a L1PA5 in the *MAP3K13* gene suggests that this chimeric insertion resulted from a recombination between two distinct RNA molecules or their simultaneous reverse transcription during retrotransposition. Another example of a chimeric insertion containing all TPRT hallmarks is shown in Figure 5D. This involves a sequence mapped to a mRNA isoform of the fragile histidine triad protein (*FHIT*) gene, an exceptionally large gene spanning 1.5 Mb of genomic DNA (Figure 5E). The upstream sequence of this LINE-1 element aligns with the first three exons of the *FHIT* gene.

**Figure 5.**
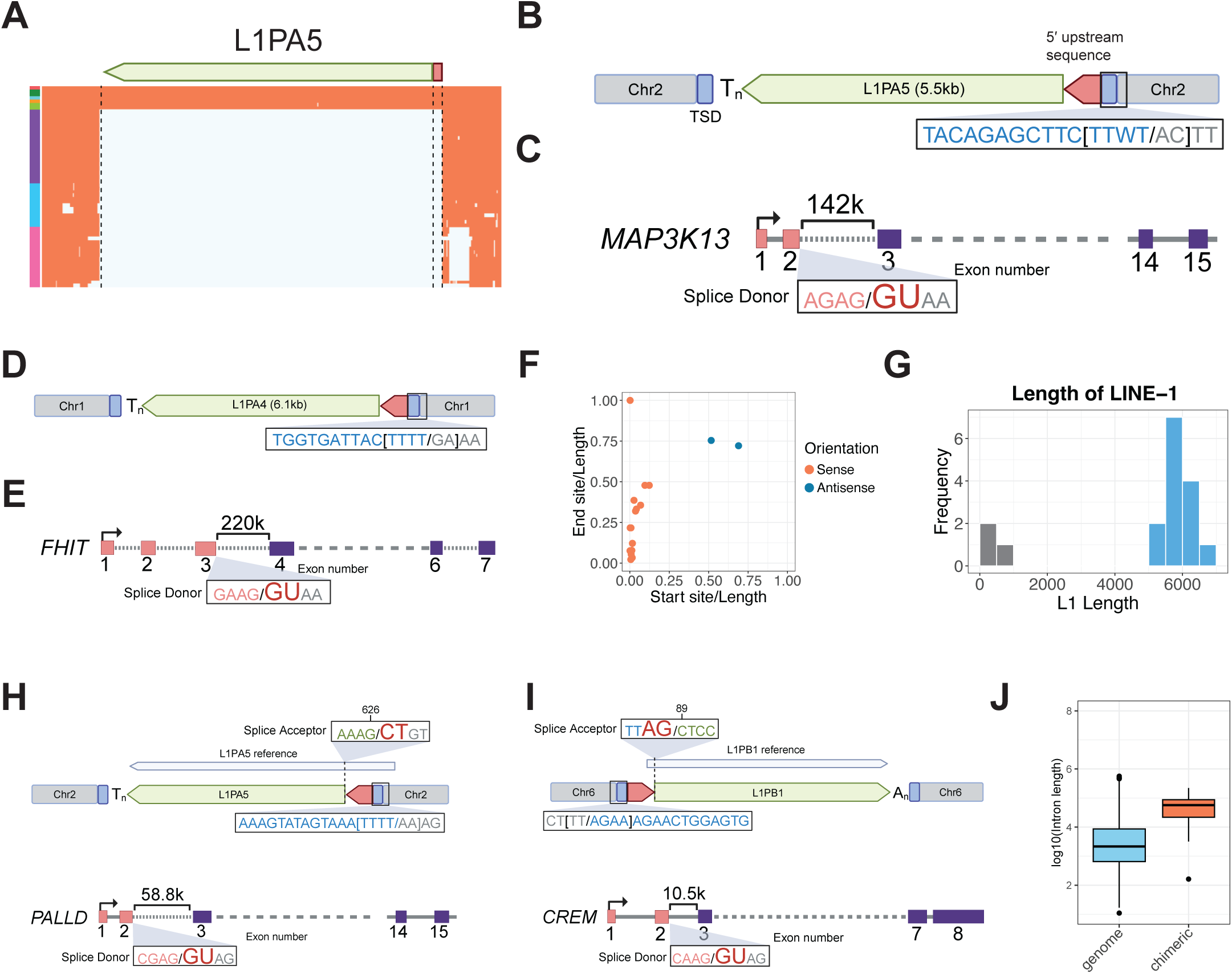
Chimeric LINE-1 insertions associated with mRNA sequences. (A) MSA showing a chimeric insertion composed of a LINE-1 element and non-LINE-1 sequence. (B) Schematic representation of the chimeric insertion structure. A 15 bp target site duplication (TSD, highlighted in blue) flanks the downstream end of the LINE-1 element and the upstream sequence. Key features include an ORF2p EN motif (in brackets) and a polythymidine sequence downstream of the LINE-1 element. (C) BLAST analysis aligns the upstream interval with the first two exons of the *MAP3K13* transcript (ENST00000424227) ending immediately at a splice donor site, diagrammed. The transcript spans 15 exons, a subset of which are diagrammed (not to scale). Distances >40 kb are shown with dashed lines. (D) Another chimeric insertion includes a 14 bp TSD, an ORF2p EN motif, and a poly(T) tail. (E) BLAST analysis of the 5′ segment aligns to the first three exons of the *FHIT* transcript (ENST00000488467), which has 7 exons spanning across 1.2 Mb. (F) A total of 17 chimeric LINE-1 insertions associated with protein-coding genes or lncRNAs were identified. Of these, 15 insertions mapped to the beginning of the transcripts, with most in the sense orientation (red) relative to LINE-1. (G) The majority of these chimeric insertions incorporate nearly full-length LINE-1 elements, longer than 5 kb (blue bars). (H) A chimeric insertion includes a 17 bp TSD, an ORF2p EN-motif, and a poly(T) tail. The upstream sequence maps to the first two exons of the *PALLD* transcript (ENST00000704822), ending immediately at a splice donor site. The LINE-1 segment starts at position 626 of an LINE-1 consensus sequence, adjacent to a previously reported splice acceptor at 625. (I) Another chimeric insertion example has all features of ORF2p-mediated retrotransposition (i.e., 16 bp TSD, an ORF2p EN motif, and a poly(A) tail). The upstream sequence maps to the first two exons of the *CREM* transcript (ENST00000354759) and terminates at the exon junction, accompanied by a splice donor, while the LINE-1 segment maps to position 89 of the L1PB1 sequence, next to a splice acceptor. (J) The median length of the introns separating the last mapped exon, and the next exon is 48 kb, significantly longer than the common first and second intron length in the genome, which has median length of 2.2kb. The differences were statistically analyzed using the Wilcoxon’s Rank Sum test, with *** denoting a p-value < 0.001.

Across all 17 of these 5′ chimeric insertions, 15 are oriented in the sense direction relative to the LINE-1 sequence. This sense orientation bias is essentially absolute in chimeric insertions wherein the 5′ piece maps to the start of a transcript and encompasses the first exon of the respective gene mRNA (Figure 5F). We observed that the first exons of some of these genes contain promoter histone marks and numerous transcription factor binding sites (TFBS) (Supplementary Figure 5), suggesting the possibility that LINE-1 can co-opt novel regulatory elements by this mechanism. Moreover, unlike chimeric LINE-1 insertions formed with repetitive RNAs (e.g., U6) or LINE-1 elements in the human genome, which frequently exhibit 5′ truncations, LINE-1 elements in these 5′ transduced chimeric structures were predominantly close to full-length (Figure 5G). This suggests that near full-length LINE-1 insertions may have a special predilection to gain extra 5′ sequence, and that the resulting chimeras may be retrotransposition-competent.

To gain insights into the mechanisms leading to these 5′ transductions, we studied the junctions between the upstream appended sequences and their associated LINE-1 element. We see that six of the 5′ transduced gene exons terminate precisely at splice donors (Figure 5C, E, H, I, Supplementary Figure 6A and Supplementary Table 4). In four cases, it was not possible to determine whether splice donors were present due to short untemplated sequences at the junction with LINE-1. In some instances, corresponding splice acceptor sites were identified within the LINE-1 sequence, including a previous known splice acceptor at position 622 of L1PA5 (76) (Figure 5H and I). However, in other cases, the corresponding splice site in LINE-1 was absent, particularly when the LINE-1 segment mapped to the first ten base pairs of the consensus sequence (Figure 5C, 5E, and Supplementary Figure 6A). In one example, a splice acceptor site was present within the LINE-1, though no corresponding splice donor site was detected (Supplementary Figure 6B). In aggregate, however, the recurring occurrence of splice sites between the 5′ exons and the downstream LINE-1 raises the possibility of *trans*-splicing involving pre-mRNAs and LINE-1 (77–81).

Potentially relevant to a *trans*-splicing model, we observe that the immediate downstream introns of the last exon included in these mRNA/LINE-1 chimeras are often longer than typical intronic intervals (Figure 5C, E, H, I, and Supplementary Figure 5A). To investigate this systematically, we compared the lengths of these downstream introns with typical lengths of the first and second introns across annotated genes in human genome. Introns associated with chimeric insertions were significantly longer overall, with a median length of 48 kb and a range of 163bp-220kb, compared to the genome-wide median of 2.2 kb (Figure 5J). These findings suggest a potential tendency for mRNAs to splice to LINE-1 sequences *in lieu* of a distant downstream splice acceptor and be captured in the genome by TPRT as 5′ segments of LINE-1 chimeric insertions.

### Co-opted promoters drive expression of 5′ compromised LINE-1

In our evaluation of 5′ transduction events, we identified two nearly full-length (5.5kb) LINE-1 insertions (Figure 6A), both missing part of the LINE-1 promoter but incorporating upstream sequences mapping to the same gene, Rap1 GTPase-GDP dissociation stimulator 1 (*RAP1GDS1*) (Figure 6B). These two insertions were found on different chromosomes and flanked by non-homologous genomic sequences and distinct TSDs (Figure 6A), indicating that the similar insertions did not the result from genomic duplication but rather independent retrotransposition events. Expanding our search for this specific *RAP1GDS1*-L1PA2 chimeric sequence, we identified a total of four insertions: one insertion on chromosome 10 and three on chromosome 1, owing to segmental duplication of one insertion on chromosome 1 (Dennis et al., 2012).

**Figure 6.**
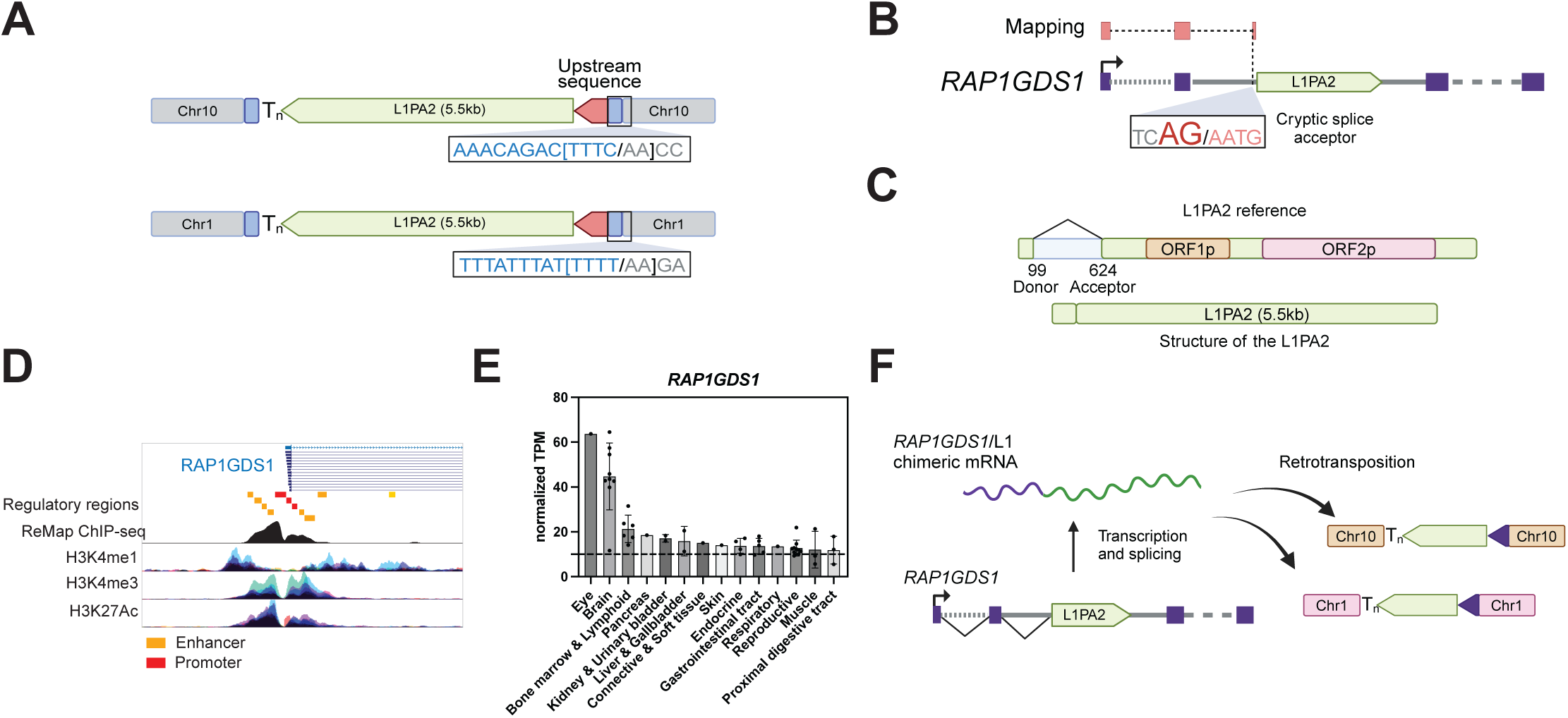
LINE-1 retrotransposition driven by a co-opted external promoter. (A) Two chimeric LINE-1 insertions, each containing highly similar LINE-1 and upstream sequences, were identified at two distinct locations. Both insertions are flanked by different TSDs of 12 bp and 13 bp and ORF2p EN motifs at their 5′ ends. (B) The upstream sequences mapped to the first two exons of *RAP1GDS1* (ENST00000508490) and a region directly upstream of a LINE-1 element located within an intron of *RAP1GDS1*. (C) A more detailed diagram of the source L1PA2 element, which is 5.5 kb in length, contains ORF1 and ORF2. Its 5′ UTR contains an interstitial deletion caused by splicing, likely rendering it transcriptionally inactivity. (D) Analysis of ChIP-seq datasets from ReMap database reveals that the *RAP1GDS1* promoter is bound by numerous transcription factors and has active histone marks, H3K4me1, H3K4me3, and H3K27Ac, suggesting high transcriptional activity. (E) GTEx and Human Protein Atlas data further supports that *RAP1GDS1* is actively expressed across multiple tissues, with particularly high expression in the brain and eye tissues. Transcripts exceeding 10 transcripts per million (TPM, marked by the horizontal line) are generally considered actively expressed. (F) A proposed mechanism for the formation of these chimeric LINE-1 insertions. Transcription of the intronic L1PA2 by the active *RAP1GDS1* promoter produces a spliced *RAP1GDS1*/L1PA2 RNA, competent for retrotransposition.

We identified a putative source L1PA2 for these insertions residing in the second intron of *RAP1GDS1* in a position to permit splicing between a donor in the second *RAP1GDS1* exon and a cryptic splice acceptor 15bp upstream of the L1PA2. While the L1PA2 has sequence corresponding to the full length of ORF1 and ORF2, it exhibits an unusual 5′ UTR, with an interstitial deletion corresponding to positions 99–624 in the L1PA2 consensus sequence (Figure 6C). This LINE-1 locus was previously identified as a Spliced Integrated Retrotransposed Element (SpIRE) and the absence of this region has been associated with the transcriptional inactivation of LINE-1 elements (76). Rather than being ‘dead-on-arrival’, which is typical of SpIREs, it appears this element co-opted the *RAP1GDS1* gene promoter to restore retrotransposition competence.

Publicly available annotations of regulatory elements in the genome indicate that the *RAP1GDS1* promoter region is highly active. According to the chromatin immunoprecipitation sequencing (ChIP-seq) database ReMap (82), this region exhibits extensive transcription factor binding across multiple cell lines. Moreover, the presence of highly enriched active histone markers, including H3K4me1, H3K4me3, and H3K27ac (Figure 6D), further supports the strong transcriptional activity of the *RAP1GDS1* promoter. Additionally, *RAP1GDS1* transcription is highly ubiquitous across tissues (Figure 6E), with particularly prominent expression in human brain. To explore this further, we analyzed *RAP1GDS1* mRNA expression in brain tissues from humans, chimpanzees, bonobos, and macaques using previously published data (55) and found high expression throughout these non-human primates (Supplementary Figure 7). We also analyzed publicly available Oxford Nanopore Technology (ONT) long-read transcriptomic sequencing data from 24 human samples, encompassing embryonic and adult tissues as well as induced pluripotent stem cells (iPSCs) (53). We found three RNA reads supporting splicing between upstream exons of *RAP1GDS1* and the intronic L1PA2: two from brain tissue and one from testis. This finding is consistent with the known high expression of *RAP1GDS1* in brain and offers evidence of the RNA intermediate likely responsible for these retrotransposition events entering the germline.

Together, these findings provide strong evidence that the *RAP1GDS1* promoter drove the expression of an intron embedded LINE-1 element, compensating its loss of intrinsic transcriptional activity and rendering it retrotransposition-competent (Figure 6F).

## DISCUSSION

### MSA is a powerful tool for TE analysis

Recent genome sequencing efforts across many species (43,83,84) are making MSA a powerful approach for identifying TE insertions, which can range in size from a few dozen base pairs to several kilobases. MSA enables the identification of polymorphic TE insertions within species (85,86), studies on TE evolution (87–89) and the discovery of new TEs (86,90). In this study, we expanded on the potential of MSA to identify chimeric LINE-1 insertions without relying on specific knowledge of those fusions *a priori*. MSA enabled us to efficiently identify and characterize insertions co-occurring at a locus around the same time. Furthermore, our use of consensus sequences from multiple genomes enhanced our annotations of key sequence features of TPRT, including the ORF2p EN cut-site motif and TSDs flanking insertions, allowing us to conclude in many instances that the sequences were indeed inserted in a single retrotransposition event. Improvements in MSA alignment tools (91) and the development of TE detection method from MSA to enhance the accurate identification and characterization of TEs (86) will no doubt continue to accelerate this field.

### Characterization of U6/LINE-1 chimeric insertions

We began this study by developing a new computational pipeline, TiMEstamp to characterize chimeric LINE-1 insertions in the human genome using MSA data. We focused first on previously known, highly abundant U6/LINE-1 chimeras and curated the largest census of U6/LINE-1 chimeric insertions to date. U6 snRNA is a critical component of the spliceosome, responsible for interacting with other snRNAs involved in the spliceosome (e.g., U2 and U4 snRNA) and interacting with the 5′ splice site of the intron in pre-mRNA (92). Consistent with previous studies, we found U6 segments in U6/LINE-1 chimeric insertions are predominantly in full-length and oriented in the sense direction with respect to the associated LINE-1. Meanwhile LINE-1 portions of these chimeras are typically 5′ truncated, consistent with reports describing RNA ligations that concatenate U6 and LINE-1 (Figure 7A) before reverse transcription (38).

**Figure 7.**
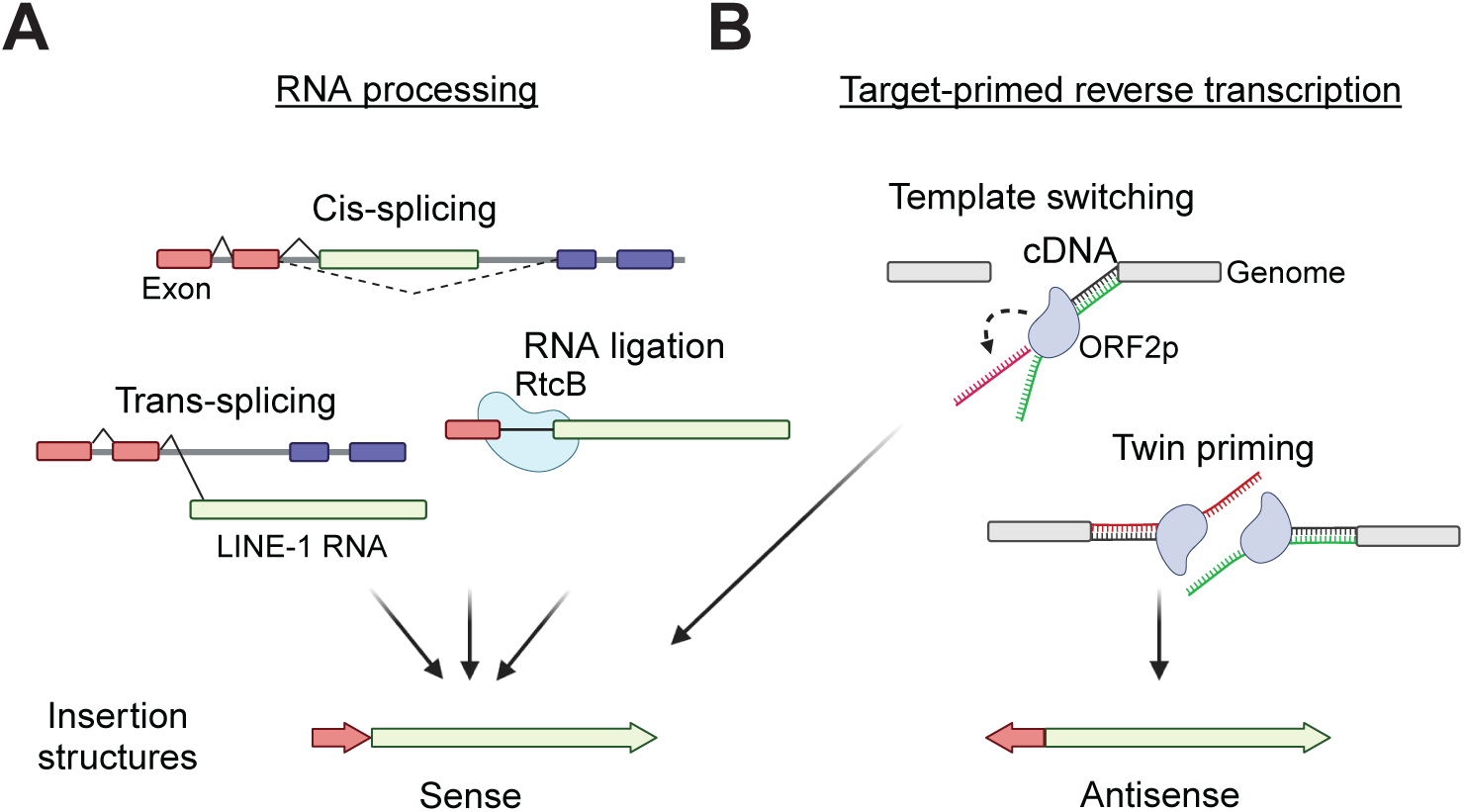
Known and proposed mechanisms for the formation of LINE-1 chimeric insertions. (A) RNA processing produces fusion transcripts that are subsequently retrotransposed. Mechanisms may include *cis* splicing, where an intronic LINE-1 is misspliced into a surrounding gene; *trans* splicing, where a LINE-1 RNA sequence is spliced with another transcript; and RNA ligation, via RtcB RNA ligase. (B) Target-primed reverse transcription (TPRT) forms the chimeric integration. This may involve template switching, where the ORF2p reverse transcriptase jumps from its primary LINE-1 RNA template to another RNA; and twin-priming, where two reverse transcription reactions are initiated on opposite ends of a double strand DNA break and joined in the resolution of the insertion. Chimeras formed by RNA processing or template switching generate insertions with non-LINE-1 and LINE-1 sequences are in the same orientation; twin-priming results chimeric segments in opposite orientations.

### Identification of new RNA repeats forming chimeric insertion with LINE-1

We expanded the species of RNAs recognized to form chimeric insertions with LINE-1 to include five new types of RNAs. Similar to previously known small, repetitive RNA that form LINE-1 chimeric insertions (33,93,94), these newly identified RNA repeats, such as Y RNA (32), tRNA (6,95), 7SL RNA (96), 7SK RNA (97,98) and 28S rRNA (99), are known to integrate into the genome as pseudogenes without forming chimeras with LINE-1 through LINE-1-mediated TPRT. Interestingly, the first reported 28 rRNA pseudogene in human was identified right next to a LINE-1 element (100), suggesting a potential association. However, the relationship between the two elements was not explored in detail at the time. Notably, previous studies have shown that U1-U6 RNA, 7SK RNA, Y RNA and 7SL RNA interact with ORF1p (98). It is possible that their association with L1 RNPs potentiates their engagement with the ORF2p reverse transcriptase.

Some of these RNA repeats have been incorporated into SINE elements in various organisms which can propagate as units by LINE machinery, for example tRNA (101–103), 7SL RNA (104,105), 5S rRNA (106–108), 28S rRNA (109) and U1/2 snRNA (110). For example, primate-specific *Alu* SINE element is derived from the deleted version of 7SL RNA before the separation of primate and Glires (104,111,112); rodent-specific B1 SINE element originated from tRNA sequence (113).

The recombination of these RNA repeats or their derived TEs with other existing retroelements can lead to the formation of novel composite mobile elements. For instance, the hominid-specific SVA element is a composite structure formed by the fusion of a LTR sequence, a variable number of tandem GC-rich repeats (VNTR), and an *Alu*-like sequence (114).

Recently, it was discovered that chimeras of RTE-type LINE and tRNA or 5S rRNA can give rise to new SINE elements (115). RTE-type LINE is a LINE subfamily that has a short 5′ UTR and only one ORF (116). This illustrates that recombination between RNA repeats creating novel sequences serves as a key driving force in genome evolution, contributing to the dynamic and rapidly evolving landscape of mobile genetic elements.

### Chimeric insertions with partial non-genetic RNAs

Our study identified many examples of LINE-1 chimeric insertions involving these so-called non-genetic RNAs. Many of these, including tRNA, 5S rRNA, 28S rRNA and U3 snoRNA have been recurrently incorporated as fragments fused to LINE-1 sequence. For tRNAs, we often see tRNA halves joined to LINE-1; it is known that cells cleave tRNAs into these half-length forms (117,118) and so it seem likely that these truncations may occur before their reverse transcription at the insertion site. tRNAs are also most commonly oriented oppositely of the associated LINE-1, suggesting they are reverse transcribed by a distinct polymerization reaction reminiscent of the twin priming model proposed for ORF2p (Figure 7B) (69). Consistent with this, previous studies have demonstrated that tRNA fragments can self-prime ORF2p-mediated retrotransposition (95). Although less is reported on processing or priming capabilities of 5S rRNA, U3 snoRNA or 28S rRNA, we see recurrent fragments of these sequences in LINE-1 chimeric insertions as well.

### *Alu* forms chimeric insertion with LINE-1

Our study also revealed that *Alu* SINE elements can form chimeric insertions with LINE-1. In these *Alu*/LINE-1 chimeras, the *Alu* portion is predominantly sense-oriented and typically full-length (∼300bp), the latter matching the general length distribution of *Alu* in the human genome. LINE-1 portions in these chimeric insertions tend to be 5′ truncated, as they are elsewhere. Like U6/LINE-1 chimeras, these features may result from RNA ligation before retrotransposition (Figure 7A) or template switching during TPRT (Figure 7B). The correlation between the ages of LINE-1 and *Alu* elements found paired together in *Alu*/LINE-1 chimeras indicates that LINE-1 most frequently forms chimeric insertions with concurrently active *Alu* species, highlighting a potential requirement for *Alu* expression or other functionality in the formation of chimeras. It is unclear if this functionality includes the ribosomal stalling capacity of *Alu* that normally permit it to co-opt ORF2p since these chimeras have a 3′ LINE-1 segment (75).

### LINE-1 may form chimeric insertions with gene mRNAs through *trans*-splicing

Leveraging our MSA-based pipeline, we identified 17 transcripts forming chimeric insertions with LINE-1. Notably, 15 of the relevant gene loci do not contain intronic LINE-1 sequences, suggesting that RNA recombination in *trans* may precede reverse transcription and genomic integration. Consistent with a model of *trans*-splicing (Figure 7A), the majority of these chimeric insertions incorporate the first exon(s) of a transcript in sense orientation with the LINE-1, and splice donor and acceptor sites are found at a proportion of the exon-LINE-1 junctions. The events appear to preferentially involve gene exons immediately upstream of long introns, consistent with prior findings that longer introns are prone to *trans* splicing (119). Interestingly, these chimeric insertions often incorporate full-length or nearly full-length LINE-1 elements, retaining the protein-coding capacity for ORF1p and ORF2p. Thus, these events frequently juxtapose the first exon of a cellular gene, sequences often enriched with TFBS, with LINE-1 protein coding sequences, potentially illustrating mechanisms for LINE-1 promoter evolution.

In contrast, the mRNA/LINE-1 chimeras with antisense-oriented mRNA did not incorporate a first exon or have features suggestive of splicing and may be more consistent with twin-priming.

### Promoter co-option by an intronic LINE-1

Two of the 17 mRNA/LINE-1 chimeras share the same internal sequence although arose through independent TPRT events. The sequence appears to have derived from a *cis* splicing event between 5′ exons of *RAP1GDS1* and an intronic LINE-1 element in that gene. This L1PA2 was previously identified as an element with an internal splice deleting approximately 500bp of its 5′ internal promoter (76) and eliminating multiple TFBS (120–124). Thus, the LINE-1 would likely not be expressed in a different genomic context. In this case, however, the element was rendered competent for retrotransposition via the activity of the *RAP1GDS1* promoter. To our knowledge, this is the first instance of a transcriptionally disabled, intronic LINE-1 co-opting an external promoter and splicing mechanisms to facilitate retrotransposition.

We speculate that this co-option of an external promoter and 5′ splicing of the *RAP1GDS1* locus (Figure 7A) may enable this element to evade LINE-1-specific silencing mechanisms. For example, the HUSH complex, which targets promoters of intronless LINE-1 elements (125,126) may not recognize intron-containing *RAP1GDS1*/L1PA2 transcripts. Binding of transcriptional suppressors, KRAB-containing zinc-finger proteins (127,128) and DNA methylation, which typically silences the LINE-1 promoter (129,130) may also be compromised by the reliance on an external promoter and the absence of 5′ LINE-1 sequence. The high and ubiquitous transcriptional activity of *RAP1GDS1* appears to have potentiated retrotransposition of this element in the germline and may have contributed to somatic mosaicism in ancestral primates until internal mutations affected the ORFs.

### Chimeric insertions may have driven LINE-1 evolution

While ORF2 and most of ORF1 sequences are relatively conserved in primates LINE-1 families, the 5′ UTR and internal promoter has been exchanged at several instances in its evolution without transitional intermediate sequences (5). We speculate that LINE-1 achieves these discrete 5′ UTR changes through its ability to recombine with other RNAs and form chimeric insertions in the genome. This may introduce entirely new binding sites for *trans* activators or permit LINE-1 to bypass host cell suppressors and explain its evolutionary success. We expect that MSA studies of chimeric and composite retroelement insertions will continue to yield insights into recombination as a driver of TE adaptability.

## DATA AVAILABILITY

The multiple sequence alignment across 470 genomes used in the analysis pipeline can be accessed from UCSC genome browser via https://hgdownload.soe.ucsc.edu/goldenPath/hg38/multiz470way/.

## AUTHOR CONTRIBUTION STATEMENT

C.-T. Law and K.H. Burns conceived the project and wrote the manuscript. C.-T. Law performed all computational analyses.

## ACKNOWLEDGMENTS

K.H. Burns and her lab are supported by NIH grants (R01CA240816, R01CA276112, R01CA289390, and UG3NS132127). C.-T. Law is supported by an American Cancer Society Postdoctoral Fellowship Award. Some figures were created using BioRender.com.

---

**Supplementary Figure 1.**
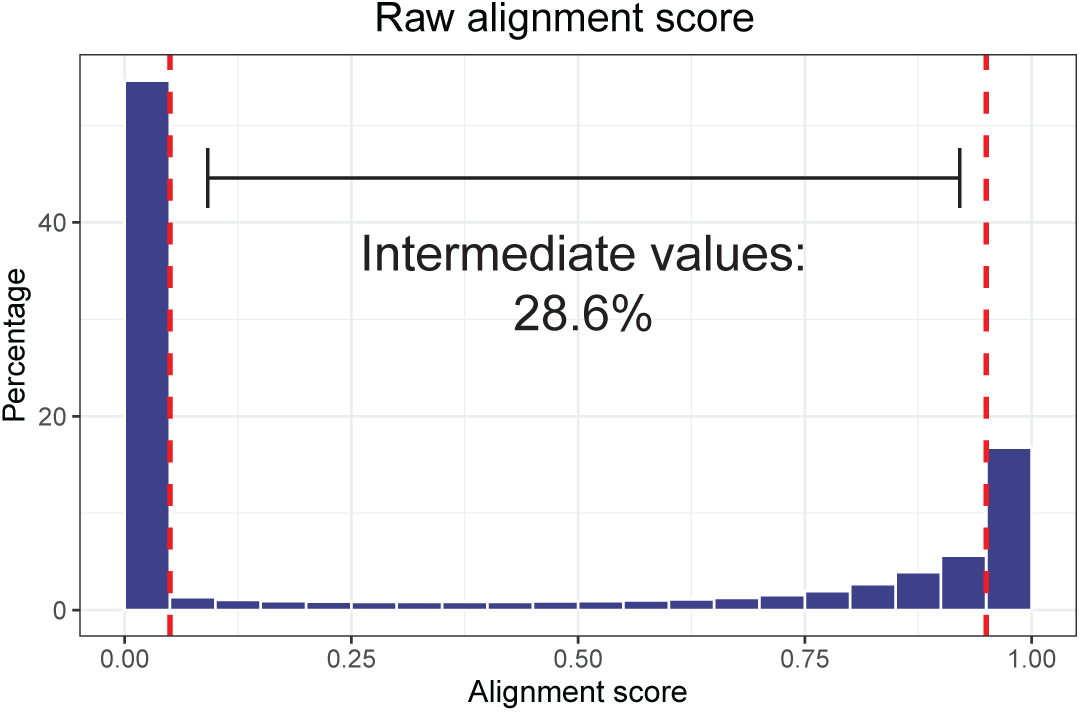
Distribution of alignment score in multiple sequence alignment across all genomes.

**Supplementary Figure 2.**
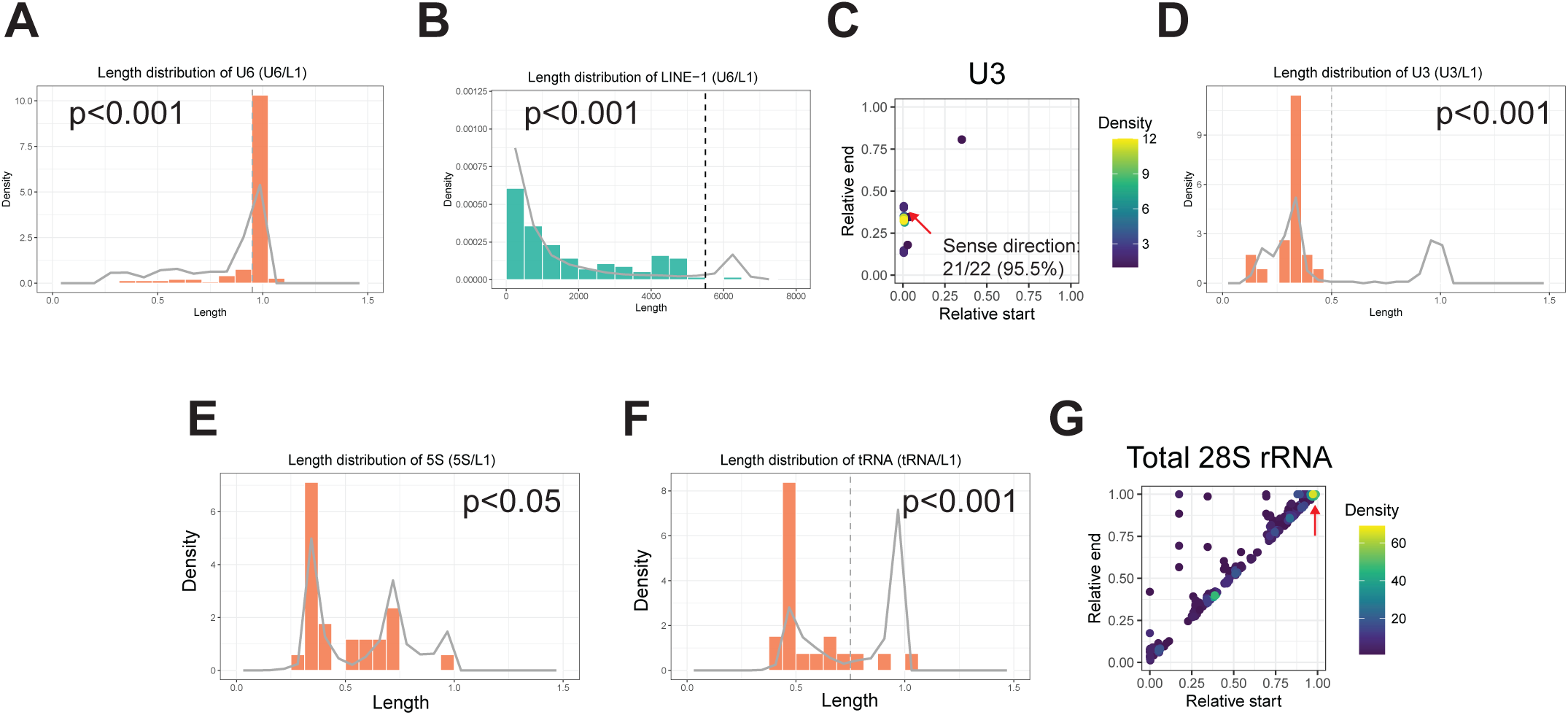
Characterization of RNA repeat associated LINE-1 chimeric insertions. (A) U6 RNA sequences in U6/LINE-1 chimeras show a significantly higher proportion of full-length sequences (defined as ≥95% of the length of the consensus U6 sequence) compared to genomic U6 RNA (indicated by the grey line), as determined by Fisher’s exact test. (B) LINE-1 sequences in U6/LINE-1 chimeras are frequently truncated (length < 5,500 bp) compared to genomic LINE-1 elements (L1PA and L1PB families). This association was statistically tested using Fisher’s exact test. (C, D) Among the identified 22 U3/LINE-1 chimeric insertions, the U3 segment is frequently truncated, with length approximately 30% of the U3 sequence. This is notably shorter when compared to U3 sequences found in the human genome. Truncation is defined as less than 50% of the full-length U3. This association was evaluated using Fisher’s exact test. Most U3 sequences are oriented in the sense direction relative to their LINE-1 counterpart. The red arrow indicates the dot with the highest data point density. (E) The length distribution of 5S rRNA in 5S/LINE-1 chimeric insertions differs significantly from those found in the genome. Due to the non-bimodal distribution of 5S RNA lengths, the statistical test used was the Wilcoxon Rank-Sum Test rather than Fisher’s exact test. (F) Compared to the abundance of full-length tRNA in the genome, chimeric LINE-1 associated tRNA are frequently truncated, where full-length tRNA is defined as 75% length of the full-length tRNA. This association was statistically evaluated using Fisher’s exact test. (G) In the genome, the insertions of 28S rRNA frequently start at the end of the 28S rRNA.

**Supplementary Figure 3.**
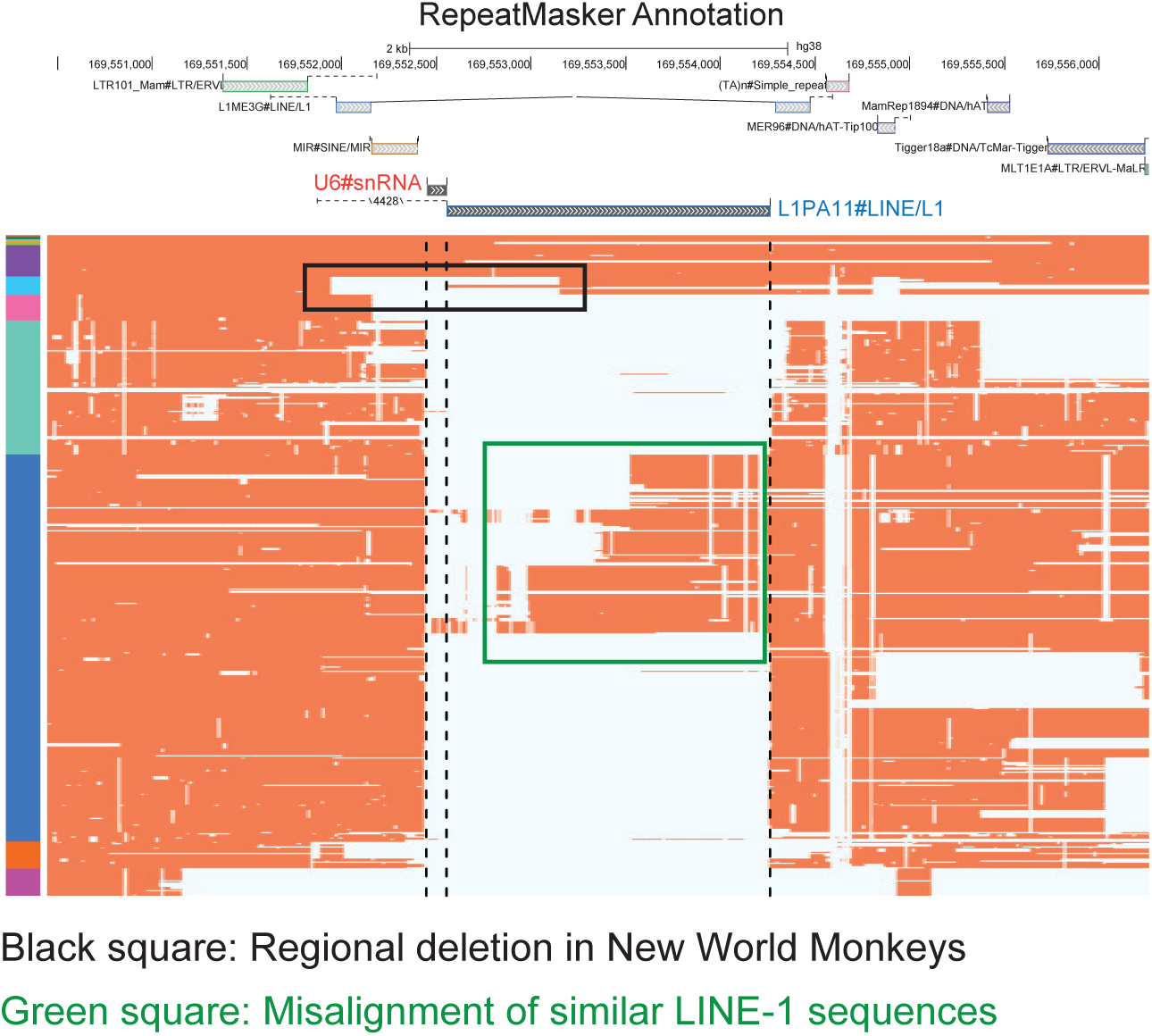
A selected example of a true U6/LINE-1 chimeric insertion filtered by the computational pipeline. The region containing the U6 sequence and part of LINE-1 (indicated by the black square) is deleted in the clade of New World monkeys. This deletion leads to an inaccurate inference of the U6 insertion timing, resulting in the exclusion of this locus due to its misclassification as a non-contemporaneous insertion. Additionally, this locus is filtered due to artifacts from multiple sequence alignment (MSA), as indicated by the green square. The portion of the first exon mapped to the chimeric insertion is highlighted in red.

**Supplementary Figure 4.**
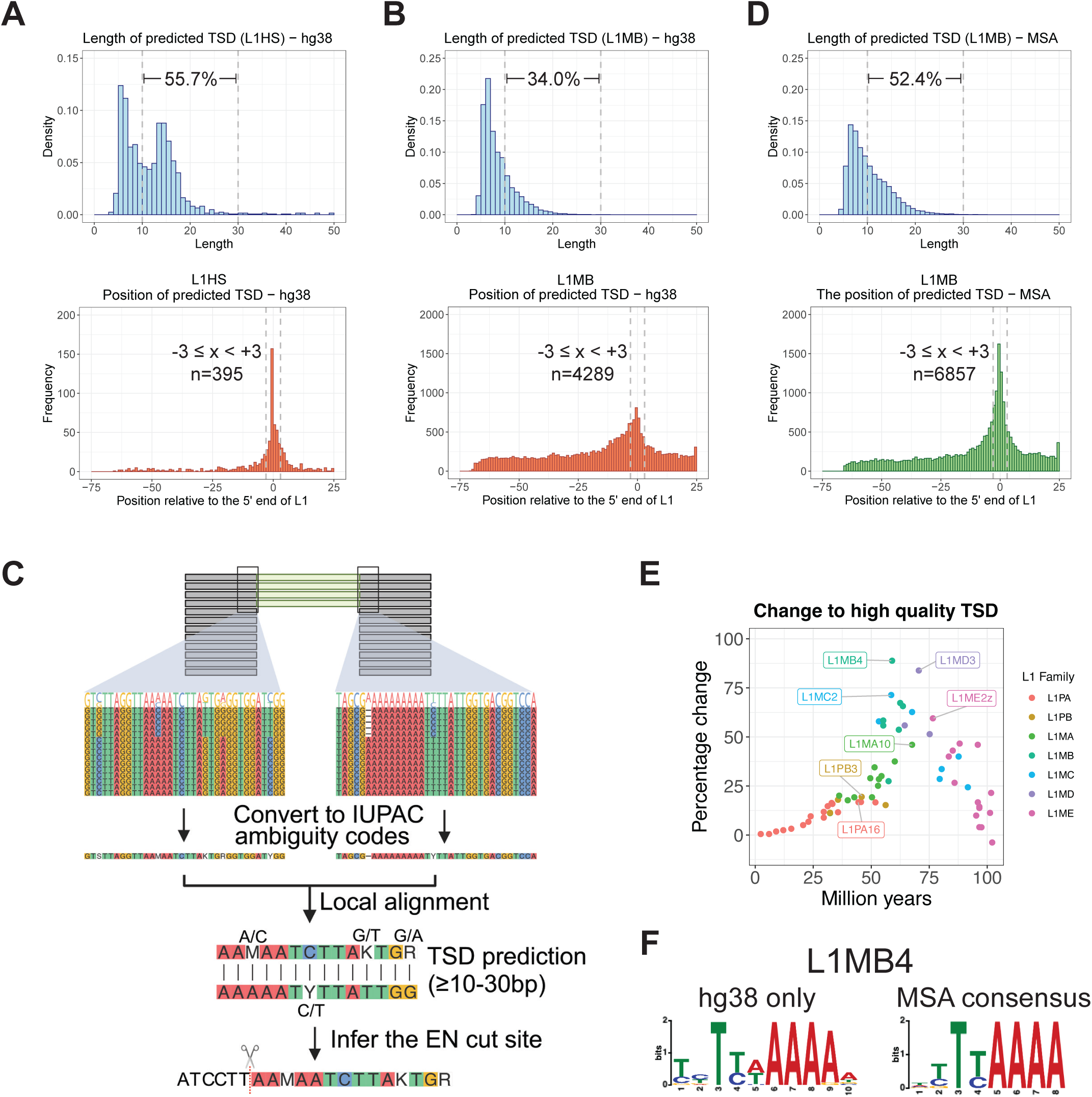
Predicting TSDs of LINE-1 from MSA (A) Upper panel: Distribution of predicted target site duplication (TSD) lengths for L1HS elements using the human reference genome hg38 only. The length of predicted TSDs shows two distinct peaks. The first peak, representing shorter lengths, indicates failed TSD predictions, while the second peak corresponds to expected TSD lengths (10-30 bp). Lower panel: Position of the predicted TSD relative to the annotated 5′ end of L1HS elements. The numbers indicate the frequency of events where the TSD is located within a 3 bp window upstream or downstream of the 5′ end of the LINE-1 insertion. (B) Upper panel: Distribution of predicted TSD lengths for L1MB elements using the human reference genome only. Lower panel: Position of the predicted TSD relative to the annotated 5′ end of L1MB elements. (C) Workflow for TSD Prediction: LINE-1 junction sequences are extracted and converted into IUPAC ambiguity codes to account for mutations in TSDs over time. These sequences are then aligned using local alignment methods to predict TSDs and EN motifs. (D) Upper panel: Distribution of predicted TSD lengths for L1MB elements using MSA cross genomes. Lower panel: Position of the predicted TSD relative to the annotated 5′ end of L1MB elements. (E) Percentage Improvement in high-quality TSD. High-quality TSDs are defined as those with a length of 10–30 bp and located within 3 bp upstream or downstream of the 5′ end of LINE-1. The percentage improvement in high-quality TSDs was calculated by comparing MSA-based predictions to predictions made exclusively using the hg38 reference genome. (F) Computed EN Motif for L1MB4 Elements: The EN motif of L1MB4 elements was predicted using MEME-suite with the sequences derived from hg38 and MSA-based approaches, demonstrating the improvement in TSD predictions for older LINE-1 families and allowing for EN motif prediction.

**Supplementary Figure 5.**
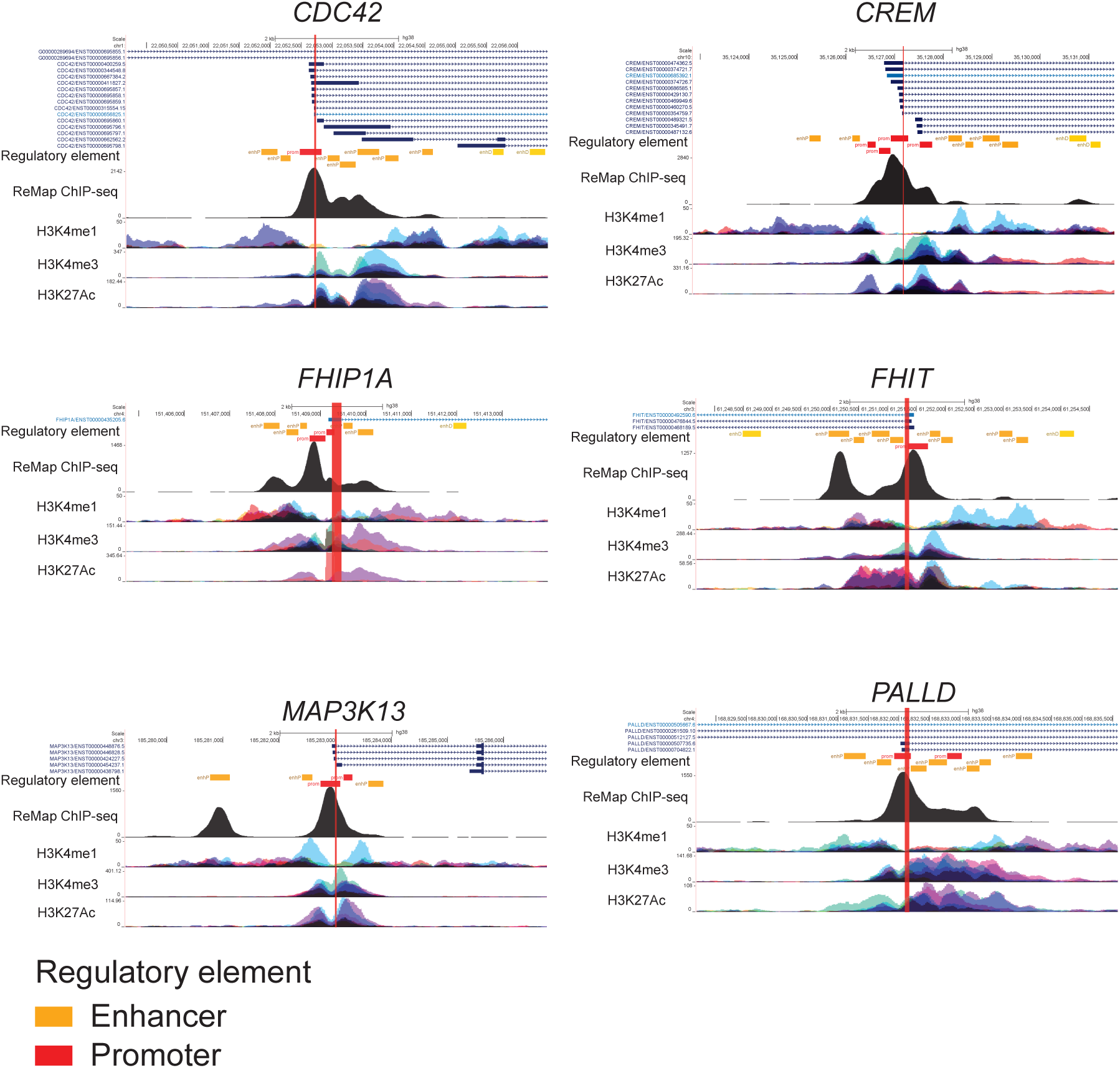
Regulatory elements overlapping with the first exon of mapped Genes. UCSC Genome Browser tracks showing regulatory elements overlapping the first exons of *CDC42*, *CREM*, *FHIP1A*, *FHIT*, *MAP3K13*, and *PALLD*. These first exons are associated with promoters enriched in transcription factor binding sites (ReMap ChIP-seq), regulatory regions and active histone marks, including H3K4me1, H3K4me3, and H3K27Ac, indicating active regulatory elements.

**Supplementary Figure 6.**
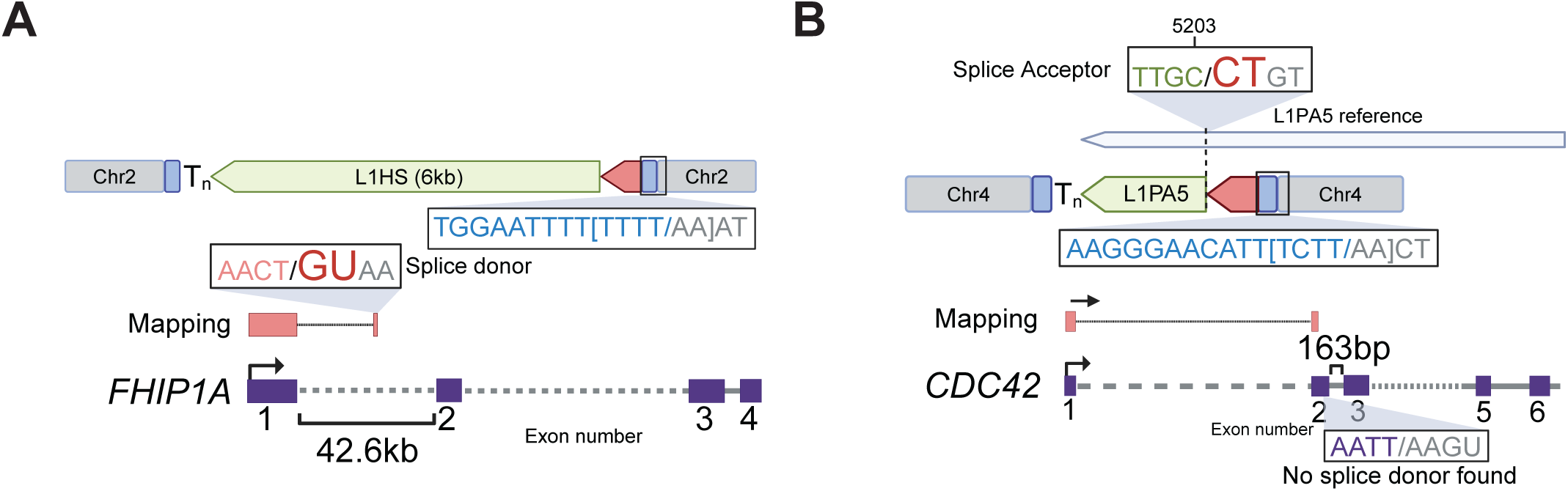
Examples of chimeric LINE-1 insertion with gene transcripts. (A) A chimeric LINE-1 insertion containing classic target site duplications (TSDs) at both ends, an endonuclease (EN) motif, and a poly(A) tail at the 3′ end. The upstream sequence of the LINE-1 element maps to an intron of *FHIP1A*, which contains a cryptic splice donor motif. The LINE-1 counterpart maps to the beginning of the L1HS reference sequence. (B) Another example of a chimeric LINE-1 insertion also contains classic TSDs at both ends, an EN motif, and a poly(A) tail at the 3′ end. The upstream sequence maps to the second exon of *CDC42*. However, no canonical splice donor motif is observed in this case. A splice acceptor motif is identified in the LINE-1 counterpart, upstream of position 5203 in a L1PA5 reference sequence.

**Supplementary Figure 7.**
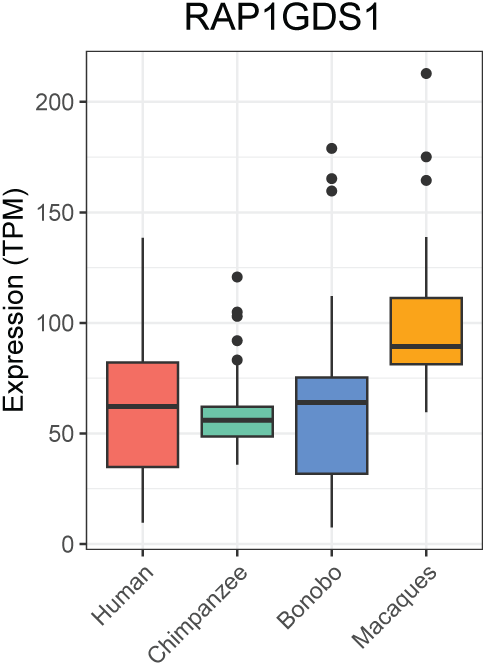
mRNA Expression levels of *RAP1GDS1* in the brain of primates.

## REFERENCE

1. Campitelli, L.F., Yellan, I., Albu, M., Barazandeh, M., Patel, Z.M., Blanchette, M. and Hughes, T.R. (2022) Reconstruction of full-length LINE-1 progenitors from ancestral genomes. Genetics, 221.

2. Sookdeo, A., Hepp, C.M., McClure, M.A. and Boissinot, S. (2013) Revisiting the evolution of mouse LINE-1 in the genomic era. Mob DNA, 4, 3.

3. Smit, A.F. (1996) The origin of interspersed repeats in the human genome. Curr Opin Genet Dev, 6, 743–748.

4. Payer, L.M. and Burns, K.H. (2019) Transposable elements in human genetic disease. Nat Rev Genet, 20, 760–772.

5. Khan, H., Smit, A. and Boissinot, S. (2006) Molecular evolution and tempo of amplification of human LINE-1 retrotransposons since the origin of primates. Genome Res, 16, 78–87.

6. Lander, E.S., Linton, L.M., Birren, B., Nusbaum, C., Zody, M.C., Baldwin, J., Devon, K., Dewar, K., Doyle, M., FitzHugh, W., et al. (2001) Initial sequencing and analysis of the human genome. Nature, 409, 860–921.

7. Kazazian, H.H., Jr. (2004) Mobile elements: drivers of genome evolution. Science, 303, 1626–1632.

8. Burton, F.H., Loeb, D.D., Voliva, C.F., Martin, S.L., Edgell, M.H. and Hutchison, C.A., 3rd. (1986) Conservation throughout mammalia and extensive protein-encoding capacity of the highly repeated DNA long interspersed sequence one. J Mol Biol, 187, 291–304.

9. Khazina, E., Truffault, V., Buttner, R., Schmidt, S., Coles, M. and Weichenrieder, O. (2011) Trimeric structure and flexibility of the L1ORF1 protein in human L1 retrotransposition. Nat Struct Mol Biol, 18, 1006–1014.

10. Khazina, E. and Weichenrieder, O. (2018) Human LINE-1 retrotransposition requires a metastable coiled coil and a positively charged N-terminus in L1ORF1p. Elife, 7.

11. Baldwin, E.T., van Eeuwen, T., Hoyos, D., Zalevsky, A., Tchesnokov, E.P., Sanchez, R., Miller, B.D., Di Stefano, L.H., Ruiz, F.X., Hancock, M., et al. (2024) Structures, functions and adaptations of the human LINE-1 ORF2 protein. Nature, 626, 194–206.

12. Thawani, A., Ariza, A.J.F., Nogales, E. and Collins, K. (2024) Template and target-site recognition by human LINE-1 in retrotransposition. Nature, 626, 186–193.

13. Weichenrieder, O., Repanas, K. and Perrakis, A. (2004) Crystal structure of the targeting endonuclease of the human LINE-1 retrotransposon. Structure, 12, 975–986.

14. Luan, D.D., Korman, M.H., Jakubczak, J.L. and Eickbush, T.H. (1993) Reverse transcription of R2Bm RNA is primed by a nick at the chromosomal target site: a mechanism for non-LTR retrotransposition. Cell, 72, 595–605.

15. Luan, D.D. and Eickbush, T.H. (1995) RNA template requirements for target DNA-primed reverse transcription by the R2 retrotransposable element. Mol Cell Biol, 15, 3882–3891.

16. Cost, G.J., Feng, Q., Jacquier, A. and Boeke, J.D. (2002) Human L1 element target-primed reverse transcription in vitro. EMBO J, 21, 5899–5910.

17. Mathias, S.L., Scott, A.F., Kazazian, H.H., Jr., Boeke, J.D. and Gabriel, A. (1991) Reverse transcriptase encoded by a human transposable element. Science, 254, 1808–1810.

18. Mendez-Dorantes, C. and Burns, K.H. (2023) LINE-1 retrotransposition and its deregulation in cancers: implications for therapeutic opportunities. Genes Dev, 37, 948–967.

19. Kazazian, H.H., Jr., Wong, C., Youssoufian, H., Scott, A.F., Phillips, D.G. and Antonarakis, S.E. (1988) Haemophilia A resulting from de novo insertion of L1 sequences represents a novel mechanism for mutation in man. Nature, 332, 164–166.

20. Holmes, S.E., Dombroski, B.A., Krebs, C.M., Boehm, C.D. and Kazazian, H.H., Jr. (1994) A new retrotransposable human L1 element from the LRE2 locus on chromosome 1q produces a chimaeric insertion. Nat Genet, 7, 143–148.

21. Moran, J.V., Holmes, S.E., Naas, T.P., DeBerardinis, R.J., Boeke, J.D. and Kazazian, H.H., Jr. (1996) High frequency retrotransposition in cultured mammalian cells. Cell, 87, 917–927.

22. Cost, G.J. and Boeke, J.D. (1998) Targeting of human retrotransposon integration is directed by the specificity of the L1 endonuclease for regions of unusual DNA structure. Biochemistry, 37, 18081–18093.

23. Morrish, T.A., Gilbert, N., Myers, J.S., Vincent, B.J., Stamato, T.D., Taccioli, G.E., Batzer, M.A. and Moran, J.V. (2002) DNA repair mediated by endonuclease-independent LINE-1 retrotransposition. Nat Genet, 31, 159–165.

24. Doucet, A.J., Wilusz, J.E., Miyoshi, T., Liu, Y. and Moran, J.V. (2015) A 3’ Poly(A) Tract Is Required for LINE-1 Retrotransposition. Mol Cell, 60, 728–741.

25. Feng, Q., Moran, J.V., Kazazian, H.H., Jr. and Boeke, J.D. (1996) Human L1 retrotransposon encodes a conserved endonuclease required for retrotransposition. Cell, 87, 905–916.

26. Dewannieux, M., Esnault, C. and Heidmann, T. (2003) LINE-mediated retrotransposition of marked Alu sequences. Nat Genet, 35, 41–48.

27. Shen, L., Wu, L.C., Sanlioglu, S., Chen, R., Mendoza, A.R., Dangel, A.W., Carroll, M.C., Zipf, W.B. and Yu, C.Y. (1994) Structure and genetics of the partially duplicated gene RP located immediately upstream of the complement C4A and the C4B genes in the HLA class III region. Molecular cloning, exon-intron structure, composite retroposon, and breakpoint of gene duplication. J Biol Chem, 269, 8466–8476.

28. Hancks, D.C., Goodier, J.L., Mandal, P.K., Cheung, L.E. and Kazazian, H.H., Jr. (2011) Retrotransposition of marked SVA elements by human L1s in cultured cells. Hum Mol Genet, 20, 3386–3400.

29. Esnault, C., Maestre, J. and Heidmann, T. (2000) Human LINE retrotransposons generate processed pseudogenes. Nat Genet, 24, 363–367.

30. Wei, W., Gilbert, N., Ooi, S.L., Lawler, J.F., Ostertag, E.M., Kazazian, H.H., Boeke, J.D. and Moran, J.V. (2001) Human L1 retrotransposition: cis preference versus trans complementation. Mol Cell Biol, 21, 1429–1439.

31. Kojima, K.K. (2010) Different integration site structures between L1 protein-mediated retrotransposition in cis and retrotransposition in trans. Mob DNA, 1, 17.

32. Perreault, J., Noel, J.F., Briere, F., Cousineau, B., Lucier, J.F., Perreault, J.P. and Boire, G. (2005) Retropseudogenes derived from the human Ro/SS-A autoantigen-associated hY RNAs. Nucleic Acids Res, 33, 2032–2041.

33. Doucet, A.J., Droc, G., Siol, O., Audoux, J. and Gilbert, N. (2015) U6 snRNA Pseudogenes: Markers of Retrotransposition Dynamics in Mammals. Mol Biol Evol, 32, 1815–1832.

34. Garcia-Perez, J.L., Doucet, A.J., Bucheton, A., Moran, J.V. and Gilbert, N. (2007) Distinct mechanisms for trans-mediated mobilization of cellular RNAs by the LINE-1 reverse transcriptase. Genome Res, 17, 602–611.

35. Buzdin, A., Ustyugova, S., Gogvadze, E., Vinogradova, T., Lebedev, Y. and Sverdlov, E. (2002) A new family of chimeric retrotranscripts formed by a full copy of U6 small nuclear RNA fused to the 3’ terminus of l1. Genomics, 80, 402–406.

36. Weber, M.J. (2006) Mammalian small nucleolar RNAs are mobile genetic elements. PLoS Genet, 2, e205.

37. Buzdin, A., Gogvadze, E., Kovalskaya, E., Volchkov, P., Ustyugova, S., Illarionova, A., Fushan, A., Vinogradova, T. and Sverdlov, E. (2003) The human genome contains many types of chimeric retrogenes generated through in vivo RNA recombination. Nucleic Acids Res, 31, 4385–4390.

38. Moldovan, J.B., Wang, Y., Shuman, S., Mills, R.E. and Moran, J.V. (2019) RNA ligation precedes the retrotransposition of U6/LINE-1 chimeric RNA. Proc Natl Acad Sci U S A, 116, 20612–20622.

39. Hecker, N. and Hiller, M. (2020) A genome alignment of 120 mammals highlights ultraconserved element variability and placenta-associated enhancers. Gigascience, 9.

40. Smit, A.F.A.H., R.; Green, P. (2013-2015) RepeatMasker Open-4.0.

41. Raney, B.J., Barber, G.P., Benet-Pages, A., Casper, J., Clawson, H., Cline, M.S., Diekhans, M., Fischer, C., Navarro Gonzalez, J., Hickey, G., et al. (2024) The UCSC Genome Browser database: 2024 update. Nucleic Acids Res, 52, D1082–D1088.

42. Hubisz, M.J., Pollard, K.S. and Siepel, A. (2011) PHAST and RPHAST: phylogenetic analysis with space/time models. Brief Bioinform, 12, 41–51.

43. Kuderna, L.F.K., Ulirsch, J.C., Rashid, S., Ameen, M., Sundaram, L., Hickey, G., Cox, A.J., Gao, H., Kumar, A., Aguet, F., et al. (2024) Identification of constrained sequence elements across 239 primate genomes. Nature, 625, 735–742.

44. Altschul, S.F., Gish, W., Miller, W., Myers, E.W. and Lipman, D.J. (1990) Basic local alignment search tool. J Mol Biol, 215, 403–410.

45. Consortium, T.R. (2019) RNAcentral: a hub of information for non-coding RNA sequences. Nucleic Acids Res, 47, D221–D229.

46. Frankish, A., Carbonell-Sala, S., Diekhans, M., Jungreis, I., Loveland, J.E., Mudge, J.M., Sisu, C., Wright, J.C., Arnan, C., Barnes, I., et al. (2023) GENCODE: reference annotation for the human and mouse genomes in 2023. Nucleic Acids Res, 51, D942–D949.

47. Ontiveros-Palacios, N., Cooke, E., Nawrocki, E.P., Triebel, S., Marz, M., Rivas, E., Griffiths-Jones, S., Petrov, A.I., Bateman, A. and Sweeney, B. (2024) Rfam 15: RNA families database in 2025. Nucleic Acids Res.

48. Quinlan, A.R. (2014) BEDTools: The Swiss-Army Tool for Genome Feature Analysis. Curr Protoc Bioinformatics, 47, 11 12 11-34.

49. Smith, T.F. and Waterman, M.S. (1981) Identification of common molecular subsequences. J Mol Biol, 147, 195–197.

50. Rice, P., Longden, I. and Bleasby, A. (2000) EMBOSS: the European Molecular Biology Open Software Suite. Trends Genet, 16, 276–277.

51. Bailey, T.L. and Elkan, C. (1994) Fitting a mixture model by expectation maximization to discover motifs in biopolymers. Proc Int Conf Intell Syst Mol Biol, 2, 28–36.

52. Jukes, T.H., Cantor, C. R. (1969) In Munro, H. N. (ed.), Mammalian Protein Metabolism. Academic Press, pp. 21–132.

53. Kaur, G., Perteghella, T., Carbonell-Sala, S., Gonzalez-Martinez, J., Hunt, T., Madry, T., Jungreis, I., Arnan, C., Lagarde, J., Borsari, B., et al. (2024) GENCODE: massively expanding the lncRNA catalog through capture long-read RNA sequencing. bioRxiv.

54. Li, H. (2018) Minimap2: pairwise alignment for nucleotide sequences. Bioinformatics, 34, 3094–3100.

55. Khrameeva, E., Kurochkin, I., Han, D., Guijarro, P., Kanton, S., Santel, M., Qian, Z., Rong, S., Mazin, P., Sabirov, M., et al. (2020) Single-cell-resolution transcriptome map of human, chimpanzee, bonobo, and macaque brains. Genome Res, 30, 776–789.

56. Hubley, R., Finn, R.D., Clements, J., Eddy, S.R., Jones, T.A., Bao, W., Smit, A.F. and Wheeler, T.J. (2016) The Dfam database of repetitive DNA families. Nucleic Acids Res, 44, D81–89.

57. Sweeney, B.A., Hoksza, D., Nawrocki, E.P., Ribas, C.E., Madeira, F., Cannone, J.J., Gutell, R., Maddala, A., Meade, C.D., Williams, L.D., et al. (2021) R2DT is a framework for predicting and visualising RNA secondary structure using templates. Nat Commun, 12, 3494.

58. Giordano, J., Ge, Y., Gelfand, Y., Abrusan, G., Benson, G. and Warburton, P.E. (2007) Evolutionary history of mammalian transposons determined by genome-wide defragmentation. PLoS Comput Biol, 3, e137.

59. Smit, A.F., Toth, G., Riggs, A.D. and Jurka, J. (1995) Ancestral, mammalian-wide subfamilies of LINE-1 repetitive sequences. J Mol Biol, 246, 401–417.

60. Harris, R.S. (2007) Ph.D. Thesis, The Pennsylvania State University, University Park, PA.

61. Blanchette, M., Kent, W.J., Riemer, C., Elnitski, L., Smit, A.F., Roskin, K.M., Baertsch, R., Rosenbloom, K., Clawson, H., Green, E.D., et al. (2004) Aligning multiple genomic sequences with the threaded blockset aligner. Genome Res, 14, 708–715.

62. Consortium, T.C.S.a.A. (2005) Initial sequence of the chimpanzee genome and comparison with the human genome. Nature, 437, 69–87.

63. Scally, A., Dutheil, J.Y., Hillier, L.W., Jordan, G.E., Goodhead, I., Herrero, J., Hobolth, A., Lappalainen, T., Mailund, T., Marques-Bonet, T., et al. (2012) Insights into hominid evolution from the gorilla genome sequence. Nature, 483, 169–175.

64. Gilbert, N., Lutz, S., Morrish, T.A. and Moran, J.V. (2005) Multiple fates of L1 retrotransposition intermediates in cultured human cells. Mol Cell Biol, 25, 7780–7795.

65. Andersen, K.L. and Collins, K. (2012) Several RNase T2 enzymes function in induced tRNA and rRNA turnover in the ciliate Tetrahymena. Mol Biol Cell, 23, 36–44.

66. Li, Z., Ender, C., Meister, G., Moore, P.S., Chang, Y. and John, B. (2012) Extensive terminal and asymmetric processing of small RNAs from rRNAs, snoRNAs, snRNAs, and tRNAs. Nucleic Acids Res, 40, 6787–6799.

67. Su, Z., Kuscu, C., Malik, A., Shibata, E. and Dutta, A. (2019) Angiogenin generates specific stress-induced tRNA halves and is not involved in tRF-3-mediated gene silencing. J Biol Chem, 294, 16930–16941.

68. Donovan, J., Rath, S., Kolet-Mandrikov, D. and Korennykh, A. (2017) Rapid RNase L-driven arrest of protein synthesis in the dsRNA response without degradation of translation machinery. RNA, 23, 1660–1671.

69. Ostertag, E.M. and Kazazian, H.H., Jr. (2001) Twin priming: a proposed mechanism for the creation of inversions in L1 retrotransposition. Genome Res, 11, 2059–2065.

70. Gilbert, N., Lutz-Prigge, S. and Moran, J.V. (2002) Genomic deletions created upon LINE-1 retrotransposition. Cell, 110, 315–325.

71. Mendez-Dorantes, C., Zeng, X., Karlow, J.A., Schofield, P., Turner, S., Kalinowski, J., Denisko, D., Lee, E.A., Burns, K.H. and Zhang, C.Z. (2024) Chromosomal rearrangements and instability caused by the LINE-1 retrotransposon. bioRxiv.

72. Zumalave, S., Santamarina, M., Espasandín, N.P., Garcia-Souto, D., Temes, J., Baker, T.M., Pequeño-Valtierra, A., Otero, I., Rodríguez-Castro, J., Oitabén, A., et al. (2024) Synchronous L1 retrotransposition events promote chromosomal crossover early in human tumorigenesis. bioRxiv, 2024.2008.2027.596794.

73. Symer, D.E., Connelly, C., Szak, S.T., Caputo, E.M., Cost, G.J., Parmigiani, G. and Boeke, J.D. (2002) Human l1 retrotransposition is associated with genetic instability in vivo. Cell, 110, 327–338.

74. Szak, S.T., Pickeral, O.K., Makalowski, W., Boguski, M.S., Landsman, D. and Boeke, J.D. (2002) Molecular archeology of L1 insertions in the human genome. Genome Biol, 3, research0052.

75. Ahl, V., Keller, H., Schmidt, S. and Weichenrieder, O. (2015) Retrotransposition and Crystal Structure of an Alu RNP in the Ribosome-Stalling Conformation. Mol Cell, 60, 715–727.

76. Larson, P.A., Moldovan, J.B., Jasti, N., Kidd, J.M., Beck, C.R. and Moran, J.V. (2018) Spliced integrated retrotransposed element (SpIRE) formation in the human genome. PLoS Biol, 16, e2003067.

77. Babiceanu, M., Qin, F., Xie, Z., Jia, Y., Lopez, K., Janus, N., Facemire, L., Kumar, S., Pang, Y., Qi, Y., et al. (2016) Recurrent chimeric fusion RNAs in non-cancer tissues and cells. Nucleic Acids Res, 44, 2859–2872.

78. Chuang, T.J., Chen, Y.J., Chen, C.Y., Mai, T.L., Wang, Y.D., Yeh, C.S., Yang, M.Y., Hsiao, Y.T., Chang, T.H., Kuo, T.C., et al. (2018) Integrative transcriptome sequencing reveals extensive alternative trans-splicing and cis-backsplicing in human cells. Nucleic Acids Res, 46, 3671–3691.

79. Yokomori, R., Kusakabe, T.G. and Nakai, K. (2024) Characterization of trans-spliced chimeric RNAs: insights into the mechanism of trans-splicing. NAR Genom Bioinform, 6, lqae067.

80. Li, H., Wang, J., Mor, G. and Sklar, J. (2008) A neoplastic gene fusion mimics trans-splicing of RNAs in normal human cells. Science, 321, 1357–1361.

81. Eul, J., Graessmann, M. and Graessmann, A. (1995) Experimental evidence for RNA trans-splicing in mammalian cells. EMBO J, 14, 3226–3235.

82. Hammal, F., de Langen, P., Bergon, A., Lopez, F. and Ballester, B. (2022) ReMap 2022: a database of Human, Mouse, Drosophila and Arabidopsis regulatory regions from an integrative analysis of DNA-binding sequencing experiments. Nucleic Acids Res, 50, D316–D325.

83. Darwin Tree of Life Project, C. (2022) Sequence locally, think globally: The Darwin Tree of Life Project. Proc Natl Acad Sci U S A, 119.

84. Zoonomia, C. (2020) A comparative genomics multitool for scientific discovery and conservation. Nature, 587, 240–245.

85. Liao, W.W., Asri, M., Ebler, J., Doerr, D., Haukness, M., Hickey, G., Lu, S., Lucas, J.K., Monlong, J., Abel, H.J., et al. (2023) A draft human pangenome reference. Nature, 617, 312–324.

86. Sierra, P. and Durbin, R. (2024) Identification of transposable element families from pangenome polymorphisms. Mob DNA, 15, 13.

87. Gibbs, R.A., Weinstock, G.M., Metzker, M.L., Muzny, D.M., Sodergren, E.J., Scherer, S., Scott, G., Steffen, D., Worley, K.C., Burch, P.E., et al. (2004) Genome sequence of the Brown Norway rat yields insights into mammalian evolution. Nature, 428, 493–521.

88. Shao, F., Zeng, M., Xu, X., Zhang, H. and Peng, Z. (2024) FishTEDB 2.0: an update fish transposable element (TE) database with new functions to facilitate TE research. Database (Oxford), 2024.

89. Osmanski, A.B., Paulat, N.S., Korstian, J., Grimshaw, J.R., Halsey, M., Sullivan, K.A.M., Moreno-Santillan, D.D., Crookshanks, C., Roberts, J., Garcia, C., et al. (2023) Insights into mammalian TE diversity through the curation of 248 genome assemblies. Science, 380, eabn1430.

90. Caspi, A. and Pachter, L. (2006) Identification of transposable elements using multiple alignments of related genomes. Genome Res, 16, 260–270.

91. Armstrong, J., Hickey, G., Diekhans, M., Fiddes, I.T., Novak, A.M., Deran, A., Fang, Q., Xie, D., Feng, S., Stiller, J., et al. (2020) Progressive Cactus is a multiple-genome aligner for the thousand-genome era. Nature, 587, 246–251.

92. Didychuk, A.L., Butcher, S.E. and Brow, D.A. (2018) The life of U6 small nuclear RNA, from cradle to grave. RNA, 24, 437–460.

93. Hasnaoui, M., Doucet, A.J., Meziane, O. and Gilbert, N. (2009) Ancient repeat sequence derived from U6 snRNA in primate genomes. Gene, 448, 139–144.

94. Jacq, C., Miller, J.R. and Brownlee, G.G. (1977) A pseudogene structure in 5S DNA of Xenopus laevis. Cell, 12, 109–120.

95. Schmitz, J., Churakov, G., Zischler, H. and Brosius, J. (2004) A novel class of mammalian-specific tailless retropseudogenes. Genome Res, 14, 1911–1915.

96. Ullu, E. and Weiner, A.M. (1984) Human genes and pseudogenes for the 7SL RNA component of signal recognition particle. EMBO J, 3, 3303–3310.

97. Murphy, S., Altruda, F., Ullu, E., Tripodi, M., Silengo, L. and Melli, M. (1984) DNA sequences complementary to human 7 SK RNA show structural similarities to the short mobile elements of the mammalian genome. J Mol Biol, 177, 575–590.

98. Goodier, J.L., Cheung, L.E. and Kazazian, H.H., Jr. (2013) Mapping the LINE1 ORF1 protein interactome reveals associated inhibitors of human retrotransposition. Nucleic Acids Res, 41, 7401–7419.

99. Wang, S., Pirtle, I.L. and Pirtle, R.M. (1997) A human 28S ribosomal RNA retropseudogene. Gene, 196, 105–111.

100. Munro, J., Burdon, R.H. and Leader, D.P. (1986) Characterization of a human orphon 28 S ribosomal DNA. Gene, 48, 65–70.

101. Daniels, G.R. and Deininger, P.L. (1985) Repeat sequence families derived from mammalian tRNA genes. Nature, 317, 819–822.

102. Okada, N. and Ohshima, K. (1993) A model for the mechanism of initial generation of short interspersed elements (SINEs). J Mol Evol, 37, 167–170.

103. Churakov, G., Smit, A.F., Brosius, J. and Schmitz, J. (2005) A novel abundant family of retroposed elements (DAS-SINEs) in the nine-banded armadillo (Dasypus novemcinctus). Mol Biol Evol, 22, 886–893.

104. Ullu, E. and Tschudi, C. (1984) Alu sequences are processed 7SL RNA genes. Nature, 312, 171–172.

105. Batzer, M.A., Deininger, P.L., Hellmann-Blumberg, U., Jurka, J., Labuda, D., Rubin, C.M., Schmid, C.W., Zietkiewicz, E. and Zuckerkandl, E. (1996) Standardized nomenclature for Alu repeats. J Mol Evol, 42, 3–6.

106. Kapitonov, V.V. and Jurka, J. (2003) A novel class of SINE elements derived from 5S rRNA. Mol Biol Evol, 20, 694–702.

107. Nishihara, H., Smit, A.F. and Okada, N. (2006) Functional noncoding sequences derived from SINEs in the mammalian genome. Genome Res, 16, 864–874.

108. Gogolevsky, K.P., Vassetzky, N.S. and Kramerov, D.A. (2009) 5S rRNA-derived and tRNA-derived SINEs in fruit bats. Genomics, 93, 494–500.

109. Longo, M.S., Brown, J.D., Zhang, C., O’Neill, M.J. and O’Neill, R.J. (2015) Identification of a recently active mammalian SINE derived from ribosomal RNA. Genome Biol Evol, 7, 775–788.

110. Kojima, K.K. (2015) A New Class of SINEs with snRNA Gene-Derived Heads. Genome Biol Evol, 7, 1702–1712.

111. Kriegs, J.O., Churakov, G., Jurka, J., Brosius, J. and Schmitz, J. (2007) Evolutionary history of 7SL RNA-derived SINEs in Supraprimates. Trends Genet, 23, 158–161.

112. Quentin, Y. (1992) Fusion of a free left Alu monomer and a free right Alu monomer at the origin of the Alu family in the primate genomes. Nucleic Acids Res, 20, 487–493.

113. Churakov, G., Sadasivuni, M.K., Rosenbloom, K.R., Huchon, D., Brosius, J. and Schmitz, J. (2010) Rodent evolution: back to the root. Mol Biol Evol, 27, 1315–1326.

114. Hancks, D.C. and Kazazian, H.H., Jr. (2010) SVA retrotransposons: Evolution and genetic instability. Semin Cancer Biol, 20, 234–245.

115. Kojima, K.K. (2018) LINEs Contribute to the Origins of Middle Bodies of SINEs besides 3’ Tails. Genome Biol Evol, 10, 370–379.

116. Malik, H.S. and Eickbush, T.H. (1998) The RTE class of non-LTR retrotransposons is widely distributed in animals and is the origin of many SINEs. Mol Biol Evol, 15, 1123–1134.

117. Lee, Y.S., Shibata, Y., Malhotra, A. and Dutta, A. (2009) A novel class of small RNAs: tRNA-derived RNA fragments (tRFs). Genes Dev, 23, 2639–2649.

118. Cole, C., Sobala, A., Lu, C., Thatcher, S.R., Bowman, A., Brown, J.W., Green, P.J., Barton, G.J. and Hutvagner, G. (2009) Filtering of deep sequencing data reveals the existence of abundant Dicer-dependent small RNAs derived from tRNAs. RNA, 15, 2147–2160.

119. Takahara, T., Tasic, B., Maniatis, T., Akanuma, H. and Yanagisawa, S. (2005) Delay in synthesis of the 3’ splice site promotes trans-splicing of the preceding 5’ splice site. Mol Cell, 18, 245–251.

120. Athanikar, J.N., Badge, R.M. and Moran, J.V. (2004) A YY1-binding site is required for accurate human LINE-1 transcription initiation. Nucleic Acids Res, 32, 3846–3855.

121. Minakami, R., Kurose, K., Etoh, K., Furuhata, Y., Hattori, M. and Sakaki, Y. (1992) Identification of an internal cis-element essential for the human L1 transcription and a nuclear factor(s) binding to the element. Nucleic Acids Res, 20, 3139–3145.

122. Becker, K.G., Swergold, G.D., Ozato, K. and Thayer, R.E. (1993) Binding of the ubiquitous nuclear transcription factor YY1 to a cis regulatory sequence in the human LINE-1 transposable element. Hum Mol Genet, 2, 1697–1702.

123. Tchenio, T., Casella, J.F. and Heidmann, T. (2000) Members of the SRY family regulate the human LINE retrotransposons. Nucleic Acids Res, 28, 411–415.

124. Yang, Z., Boffelli, D., Boonmark, N., Schwartz, K. and Lawn, R. (1998) Apolipoprotein(a) gene enhancer resides within a LINE element. J Biol Chem, 273, 891–897.

125. Seczynska, M., Bloor, S., Cuesta, S.M. and Lehner, P.J. (2022) Genome surveillance by HUSH-mediated silencing of intronless mobile elements. Nature, 601, 440–445.

126. Liu, N., Lee, C.H., Swigut, T., Grow, E., Gu, B., Bassik, M.C. and Wysocka, J. (2018) Selective silencing of euchromatic L1s revealed by genome-wide screens for L1 regulators. Nature, 553, 228–232.

127. de Tribolet-Hardy, J., Thorball, C.W., Forey, R., Planet, E., Duc, J., Coudray, A., Khubieh, B., Offner, S., Pulver, C., Fellay, J., et al. (2023) Genetic features and genomic targets of human KRAB-zinc finger proteins. Genome Res, 33, 1409–1423.

128. Playfoot, C.J., Duc, J., Sheppard, S., Dind, S., Coudray, A., Planet, E. and Trono, D. (2021) Transposable elements and their KZFP controllers are drivers of transcriptional innovation in the developing human brain. Genome Res, 31, 1531–1545.

129. Ewing, A.D., Smits, N., Sanchez-Luque, F.J., Faivre, J., Brennan, P.M., Richardson, S.R., Cheetham, S.W. and Faulkner, G.J. (2020) Nanopore Sequencing Enables Comprehensive Transposable Element Epigenomic Profiling. Mol Cell, 80, 915–928 e915.

130. Sanchez-Luque, F.J., Kempen, M.H.C., Gerdes, P., Vargas-Landin, D.B., Richardson, S.R., Troskie, R.L., Jesuadian, J.S., Cheetham, S.W., Carreira, P.E., Salvador-Palomeque, C., et al. (2019) LINE-1 Evasion of Epigenetic Repression in Humans. Mol Cell, 75, 590–604 e512.

